# An open imaging and AI resource enabling unbiased quantification of extrachromosomal DNA at scale

**DOI:** 10.64898/2026.09.18.752721

**Authors:** Poorya Behnamie, River Summers, Jingting Chen, Aarav Mehta, Adesuwa Igbinigie, Qin Liu, Molly Murray, Damien Guilbaud, Marc Niethammer, Elizabeth Brunk

**Affiliations:** Integrative Program for Biological and Genome Sciences (IBGS), University of North Carolina at Chapel Hill, Chapel Hill, NC 27516; Department of Computer Science, University of North Carolina at Chapel Hill, Chapel Hill, NC 27516; Department of Biochemistry and Biophysics, University of North Carolina at Chapel Hill, Chapel Hill, NC 27516; Department of Computer Science and Engineering, and Department of Neurological Surgery, University of California San Diego; Department of Chemistry, University of North Carolina at Chapel Hill, Chapel Hill, NC 27516; Department of Pharmacology, University of North Carolina at Chapel Hill, Chapel Hill, NC 27516; Computational Medicine Program, University of North Carolina at Chapel Hill, Chapel Hill, NC 27516; Lineberger Comprehensive Cancer Center, University of North Carolina at Chapel Hill, Chapel Hill, NC 27516

**Author notes:** Correspondence should be addressed to: Elizabeth Brunk.

## Abstract

Quantitative imaging of extrachromosomal DNA (ecDNA) is increasingly important for studying cancer heterogeneity and adaptation, yet automated analysis has been limited by the absence of accessible imaging data, gold standard annotations and adaptable computational tools. Here we establish an open resource for computational ecDNA imaging, integrating 2,986 native-resolution metaphase FISH image sets with manual annotations, standardized benchmarks and open-source quantification frameworks. We use this resource to systematically compare existing and newly developed approaches spanning rule-based computer vision, deep-learning-based segmentation and probabilistic localization. This comparison reveals count-dependent underestimation that distorts ecDNA copy-number distributions and motivates ecCount, a probabilistic localization method developed here to preserve individual ecDNA signals and quantitative burden. ecCount achieves an object-level F1 score of 0.939 on held-out images with minimal count bias. Together, the images, annotations, retrainable models, evaluation tools and guided workflows provide community infrastructure for applying, adapting and improving automated ecDNA quantification across experimental systems.

---

Extrachromosomal DNA (ecDNA) has emerged as an important driver of tumor evolution^1–3^, cellular heterogeneity^4–6^ and treatment resistance^2,7^. Because ecDNA lacks centromeres and is inherited unequally during cell division^8–10^, genetically similar cells can differ markedly in ecDNA-based oncogene copy numbers^11,12^. The resulting distribution of ecDNA across individual cells can shape transcription^13,14^, fitness and adaptation^12,15,16^ under selective pressure. Measuring this variation directly is therefore central to understanding how ecDNA contributes to cancer evolution.

Metaphase fluorescence in situ hybridization (FISH) remains one of the only approaches that directly quantifies ecDNA in individual cells and distinguishes extrachromosomal from chromosome-associated amplification. Yet its quantitative analysis remains largely manual. Conventional cytogenetic workflows^17,18^ typically examine tens of metaphases, sufficient to identify chromosome abnormalities but too few to resolve the distribution of ecDNA states across a heterogeneous cell population. Individual cells within the same population can harbor fewer than ten to more than one thousand ecDNA copies, and the prevalence of these states can shift during treatment^19,20^, selection and adaptation^15,16^. Measuring these dynamics therefore requires moving from tens to hundreds or thousands of cells. At this scale, image analysis and manual annotation become major bottlenecks, creating a need for computational approaches that can reliably identify and count individual ecDNA signals across heterogeneous FISH images.

Computational approaches have begun to automate ecDNA detection^4,19,21^, but progress has been constrained by the absence of shared imaging resources and common evaluation standards. Images and annotations have largely remained within individual studies, while existing models are often difficult to retrain or adapt to new experimental settings. Methods have also been developed using different image collections, annotation strategies and output formats, making it difficult to determine how they compare, where they fail or whether advances in computer vision improve the biological measurement itself. This lack of open infrastructure is a major limitation for a rapidly expanding field in which quantitative imaging will increasingly be needed to measure ecDNA heterogeneity and dynamics at scale.

Here, we establish a reference resource for computational analysis of ecDNA in metaphase FISH images. The resource comprises nearly 3,000 native-resolution image sets across five ecDNA-positive cancer cell lines, including a 1,145-image manually annotated benchmark with fixed training, validation and held-out test partitions. Using this common framework, we compare rule-based object detection, deep-learning-based segmentation and probabilistic localization, and introduce ecCount, which reformulates ecDNA detection around the probable location of individual signals rather than fixed object boundaries. Beyond benchmarking these approaches, we release the imaging data, annotations, model outputs, evaluation code and open-source implementations together with interactive Jupyter notebook tutorials that guide users through preparing their own data, retraining the models and evaluating performance within the same framework. This resource is therefore designed not only to benchmark current ecDNA image-analysis methods, but to provide a practical foundation that the community can use, adapt and extend as new datasets and computational approaches emerge.

## A reference resource enables comparison of distinct ecDNA quantification paradigms

Quantifying ecDNA from metaphase FISH images requires distinguishing chromosomes from extrachromosomal fluorescent signals and counting each ecDNA object within an individual cell. Automation is difficult because both image appearance and ecDNA biology are highly variable. Probe intensity and background differ between images, chromosomes vary in size, shape and arrangement, and amplified loci can appear either on chromosomes or as extrachromosomal signals (**Fig. 1e**).

**Figure 1:**
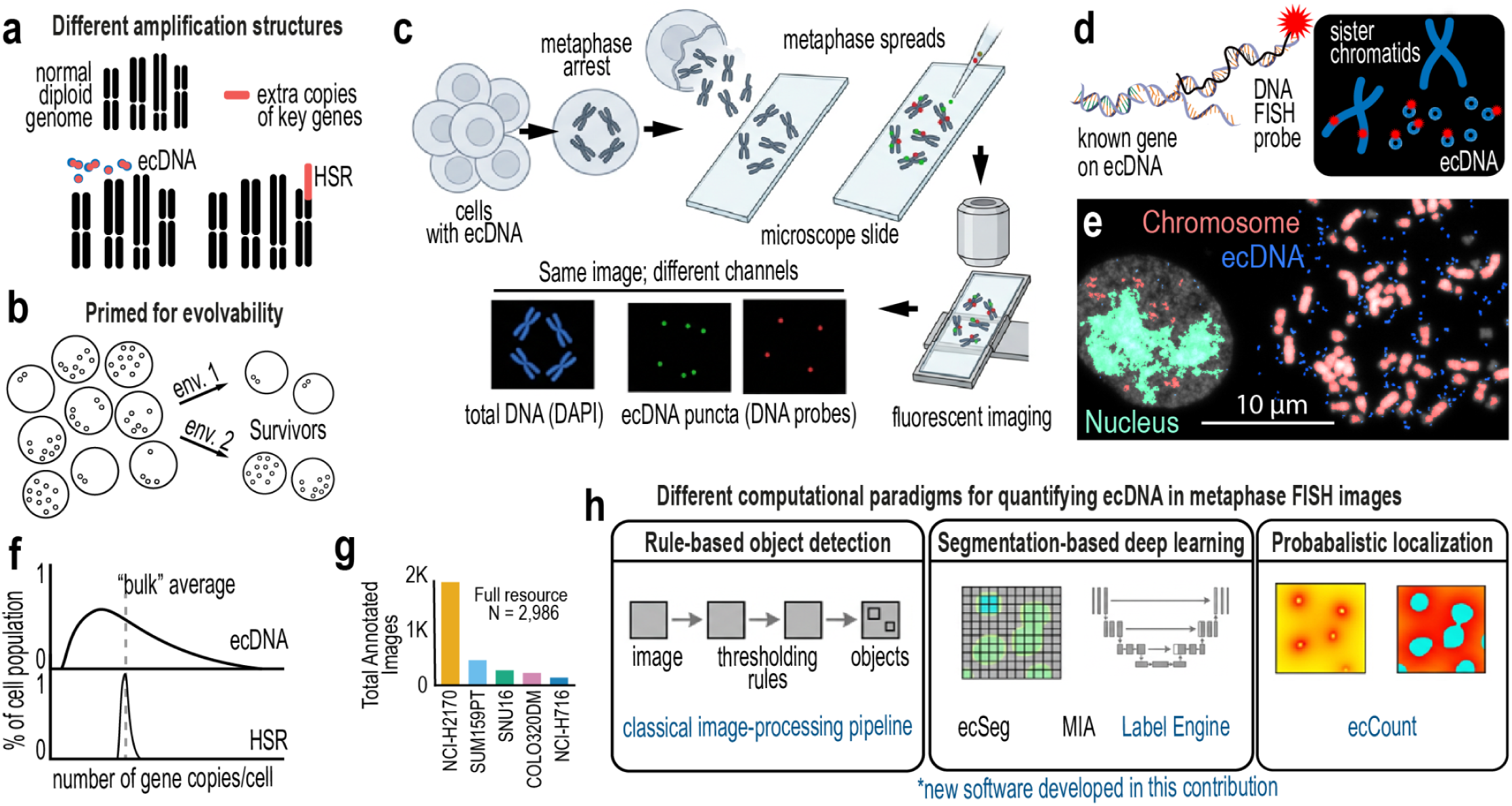
A reference imaging resource captures the complexity of ecDNA detection. a, Schematic comparison of gene amplifications residing in extrachromosomal DNA (ecDNA) and chromosome-associated amplification as homogeneously staining regions (HSRs). **b,** Cell-to-cell variation in ecDNA abundance generates heterogeneous states that can support adaptation, tumor evolution and treatment resistance. **c,** Metaphase FISH workflow for direct visualization of ecDNA. **d,** Locus-specific fluorescent probes identify amplified sequences on chromosomes and as extrachromosomal signals. **e,** Representative metaphase FISH images illustrating the classification problem and variation in chromosome organization, fluorescence intensity, background and ecDNA abundance. **f,** Distribution of ecDNA and HSR abundance across individual cells, illustrating the broad cell-to-cell variation in ecDNA copy number. **g,** Composition of the reference imaging resource, comprising 2,986 paired probe-channel RGB and DAPI image sets from five ecDNA-positive cancer cell lines, with manual ecDNA annotations and manually defined regions of interest (ROI) for the benchmark subset (n=1,145). **h,** Three computational paradigms evaluated for automated ecDNA quantification: rule-based object detection (classical pipeline), deep-learning-based segmentation (ecSeg, MIA, Label Engine) and probabilistic localization (ecCount). The classical pipeline, Label Engine and ecCount were fitted to the benchmark training partition; MIA predictions are archived outputs of the original study’s model, and ecSeg was applied zero-shot with its released weights.

The abundance of ecDNA is similarly heterogeneous: cells within the same population can harbor fewer than ten to more than one thousand ecDNA copies, whereas chromosome-based amplifications, or homogeneously staining regions (HSRs^22,23^), typically span a narrower range (**Fig. 1f**). Dense ecDNA regions contain closely spaced or overlapping signals that are difficult to resolve individually, whereas in sparse cells even a few errors can distort the measured count. Manual annotation adds further uncertainty because annotators may mark object centers, boundaries or larger structures, and placement can vary for small or overlapping signals^19,24^. Thus, computational methods must learn from annotations that contain both biological ambiguity and spatial uncertainty.

To provide a common foundation for method development, we assembled 2,986 native-resolution metaphase FISH image sets from five ecDNA-positive cancer cell lines^6,25–27^ spanning different amplified loci and ecDNA burdens (**Fig. 1g**). Each set contains matched probe-channel RGB and DAPI (4′,6-diamidino-2-phenylindole) images with ecDNA annotations. We evaluated three computational paradigms: rule-based object detection, deep-learning-based segmentation and probabilistic localization (**Fig. 1h**). Rule-based approaches identify objects from image properties such as intensity, size and shape^28,29^; segmentation models learn which pixels correspond to ecDNA; and probabilistic localization models learn where individual signals are likely to occur.

For direct comparison, we established a 1,145-image benchmark from four cell lines with manually defined metaphase regions of interest (ROIs) and fixed training (800 images), validation (170) and held-out test (175) partitions (**Fig. 2a,b; *Extended Data Fig. 1***). Predicted ROIs are additionally provided across the resource, while all primary benchmark analyses use manually defined regions. ROIs excluded unburst nuclei, neighboring cells and features outside the quantified metaphase (**Fig. 2c*; Extended Data Fig. 2***), and individual ecDNA signals were annotated as point coordinates (**Fig. 2d**).

**Figure 2:**
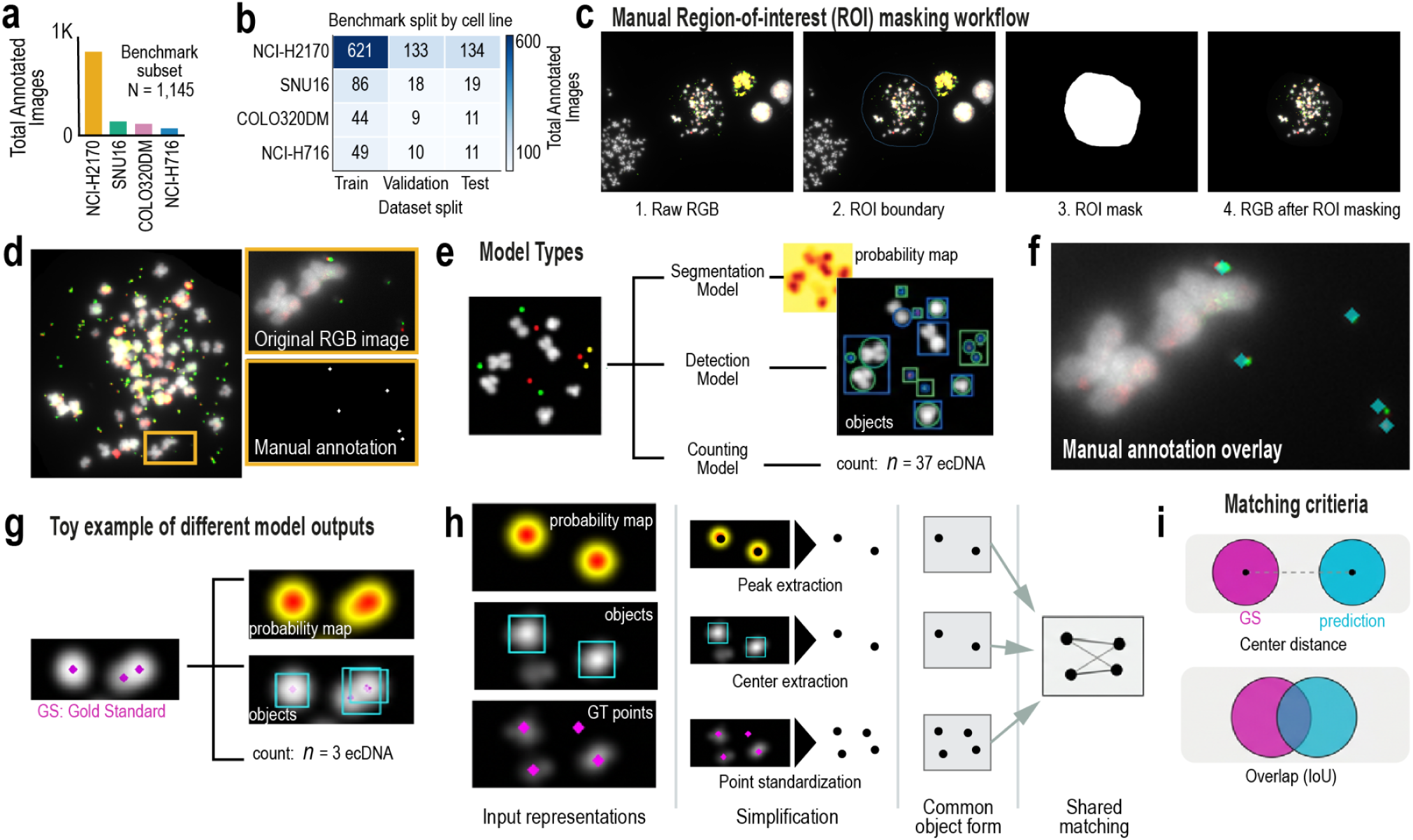
A unified benchmark enables comparison of distinct ecDNA quantification paradigms. a, Composition of the 1,145-image ROI-constrained benchmark across four ecDNA-positive cancer cell lines. Images contain matched DAPI and probe-channel RGB data with one or two locus-specific FISH probes. **b,** Distribution of benchmark images across cell lines and fixed training, validation and held-out test partitions. **c,** Manual region-of-interest annotation used to isolate individual metaphase spreads and exclude unburst nuclei, neighboring cells and other image features outside the analyzed cell. **d,** Manual ecDNA annotation (gold standard): one point per fluorescent ecDNA signal, placed at the approximate center of the signal. **e,** Computational approaches produce different forms of prediction, including pixel-wise probability maps, binary masks and discrete object locations. **f,** Illustration of uncertainty in manual annotation. Small differences in annotation placement can identify the same biological signal while producing different pixel-level masks. **g,** Representative outputs generated by segmentation, object-detection and localization approaches. **h,** Unified evaluation framework. Heterogeneous model outputs are converted to a common object-level representation and compared with manual annotations using spatially tolerant matching, enabling performance to be quantified by object detection, pixel localization and ecDNA count. **i,** Matching criteria. A prediction and a gold-standard object are eligible for matching if the centroid distance is at most 20 pixels or the mask intersection over union (IoU) is at least 0.1 (OR policy); eligible pairs are then assigned one-to-one at minimum combined distance–overlap cost.

The three paradigms generate different outputs: object bounding boxes, binary masks and probability maps (**Fig. 2e**). Because manual annotation is itself spatially uncertain^24^, small differences in annotation placement can identify the same biological signal while differing at the pixel level (**Fig. 2f**). We converted every output to connected components of a binary mask (**Fig. 2g,h**), rendering point outputs and annotations as 5 × 5-pixel diamonds. Predicted objects were then matched one-to-one to annotated objects within a distance or overlap tolerance^28^ (**Fig. 2i*; Extended Data Fig. 3***), and performance was quantified as object-level F1 (from matched and unmatched objects pooled across images), pixel-level Dice and per-image count error.

## Optimized classical computer vision provides a few-parameter baseline

Classical computer vision uses annotations only to tune a few parameters, not to learn image features. We developed a rule-based pipeline in which ROI-restricted probe-channel images undergo signal enhancement, thresholding and connected-component analysis^30^, followed by filtering and classification of ecDNA-like and chromosome-associated objects (**Fig. 3a-b**).

**Figure 3:**
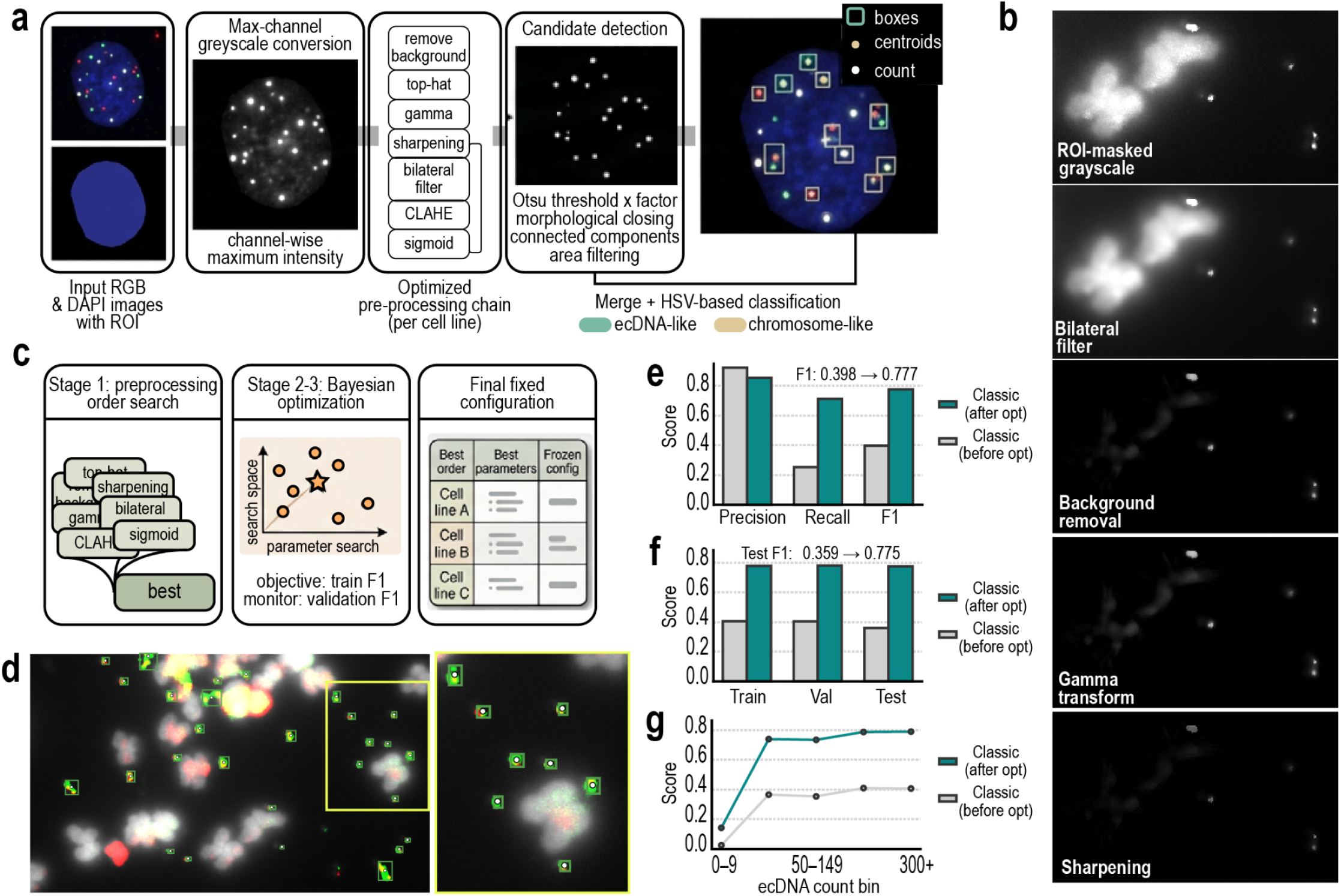
Optimization enables low-supervision ecDNA detection by classical computer vision. a, Classical computer-vision pipeline. ROI-restricted probe images undergo channel-wise intensity transformation and optimized preprocessing, followed by thresholding, morphological processing, connected-component detection, area filtering and HSV-based classification of ecDNA-like and chromosome-like objects. Detected ecDNA objects are represented by bounding boxes and centroids and converted to image-level counts. **b,** Representative images showing progressive transformation of the FISH signal through the preprocessing and detection pipeline. **c,** Parameter-optimization strategy. Preprocessing order was first selected by systematic search, followed by Bayesian optimization of preprocessing and detection parameters using training performance and validation F1 for model selection. Final configurations were optimized independently by cell line and frozen before evaluation on the held-out test set. **d,** Representative held-out metaphase showing predicted ecDNA objects, with an enlarged region illustrating individual detections. **e,** Object-level precision, recall and F1 before and after parameter optimization. **f,** Object-level F1 across fixed training, validation and held-out test partitions before and after optimization. **g,** Object-level F1 by gold-standard ecDNA count per image (count bins). Performance was lowest in low-burden images and increased across higher ecDNA-count bins; optimization reduced count mean absolute error (MAE) from 144.2 to 44.9 ecDNA per image.

We optimized preprocessing order and detection settings^31^ for each cell line on training images and froze the configuration with the highest validation F1; test images were not used (**Fig. 3c*; Extended Data Fig. 4a-d***). The optimized pipeline recovered substantially more ecDNA signals but continued to merge neighboring objects in crowded regions (**Fig. 3d**). Object-level F1 increased from 0.398 to 0.777 over all images (**Fig. 3e*; Extended Data Fig. 4e-f***), and from 0.359 to 0.775 on held-out test images (**Fig. 3f**).

Performance was least stable in low-burden images, while optimization markedly reduced count error (**Fig. 3g**). Thus, classical vision achieves substantial accuracy with few tuned parameters, but context-specific parameterization and connected-component detection constrain performance in heterogeneous images.

## Deep-learning-based segmentation undercounts crowded ecDNA signals

Deep-learning approaches commonly formulate ecDNA detection as semantic segmentation^29,32^, in which manual annotations are converted into pixel-level masks used to train a network to distinguish ecDNA from background (**Fig. 4a**). We evaluated this paradigm with Label Engine, a U-Net developed here and trained on the benchmark training partition (**Fig. 4b**), and with two published models that were not retrained: MIA^19^, from archived predictions of the original study’s model, and ecSeg^21^, applied zero-shot because our ecDNA-only annotations cannot supervise its four-class output (Methods).

**Figure 4:**
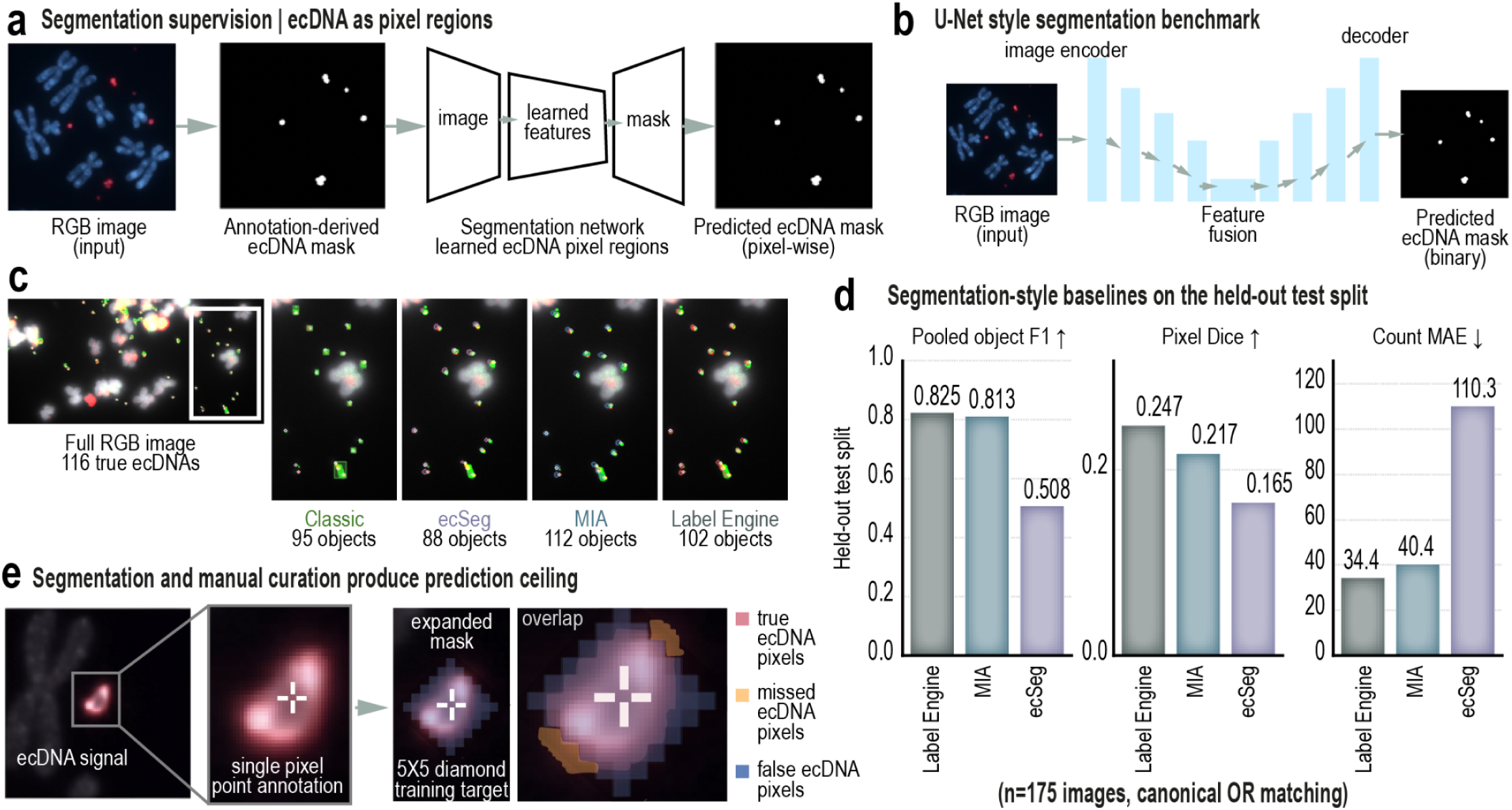
Segmentation-based learning undercounts crowded ecDNA signals. a, Segmentation-based formulation of ecDNA detection. Manual annotations are converted into pixel-defined ecDNA regions used to train a network to distinguish ecDNA foreground from background. **b,** Label Engine segmentation network: a residual convolutional image encoder and a U-Net decoder with skip connections, producing per-pixel background and ecDNA probabilities. The network was run in its automatic mode, which takes the image alone; no prompts were used for training or inference. **c,** Representative metaphase with 116 annotated ecDNAs (gold standard) and predictions from the optimized classical pipeline, ecSeg, MIA and Label Engine. Predicted object counts are 95, 88, 112 and 102, respectively; enlarged regions illustrate differences in recovery of individual fluorescent signals. **d,** Performance of Label Engine, MIA and ecSeg on the 175-image held-out test set: object-level F1, pixel-level Dice and count mean absolute error. Label Engine was trained on the benchmark training partition, MIA predictions are archived outputs of the original study’s model, and ecSeg was applied zero-shot with its released weights. Dice is low for all methods because the gold-standard masks are 5 × 5-pixel diamonds rather than object outlines (Methods). Object-level F1 was 0.825 for Label Engine, 0.813 for MIA and 0.508 for ecSeg. **e,** Schematic illustrating the limitation of hard-mask supervision. Point annotations are converted into fixed foreground pixels (5 × 5-pixel diamonds), and neighboring ecDNA signals can merge into continuous regions that preserve pixel overlap while reducing the number of discrete objects recovered.

All three approaches recovered visible ecDNA signals but differed markedly in their ability to preserve individual objects (**Fig. 4c**). On the held-out test set, object-level F1 was 0.825 for Label Engine, 0.813 for MIA and 0.508 for ecSeg (zero-shot), with corresponding differences in pixel-level Dice and count error (**Fig. 4d**).

A key limitation arises from how segmentation training labels are constructed. Manual annotation identifies the presence of an ecDNA signal but does not define its exact physical boundary. Training therefore requires converting uncertain annotations into fixed foreground regions (**Fig. 4e**). When neighboring annotations or predictions overlap, several ecDNA signals can collapse into a single connected region^24,33,34^ even when the model correctly identifies the broader area containing ecDNA.

Segmentation models can therefore place foreground in approximately the correct location while merging adjacent ecDNAs and underestimating their number, revealing a mismatch between pixel segmentation and a biological measurement defined by discrete object number and position.

## Probabilistic localization preserves individual ecDNA signals

We therefore reformulated ecDNA detection as a localization problem: rather than asking which pixels belong to ecDNA, ecCount asks where individual ecDNA signals are most likely to occur. A U-Net^32,35^ transforms each FISH image into a continuous probability map in which local maxima represent likely ecDNA positions (**Fig. 5a**).

**Figure 5:**
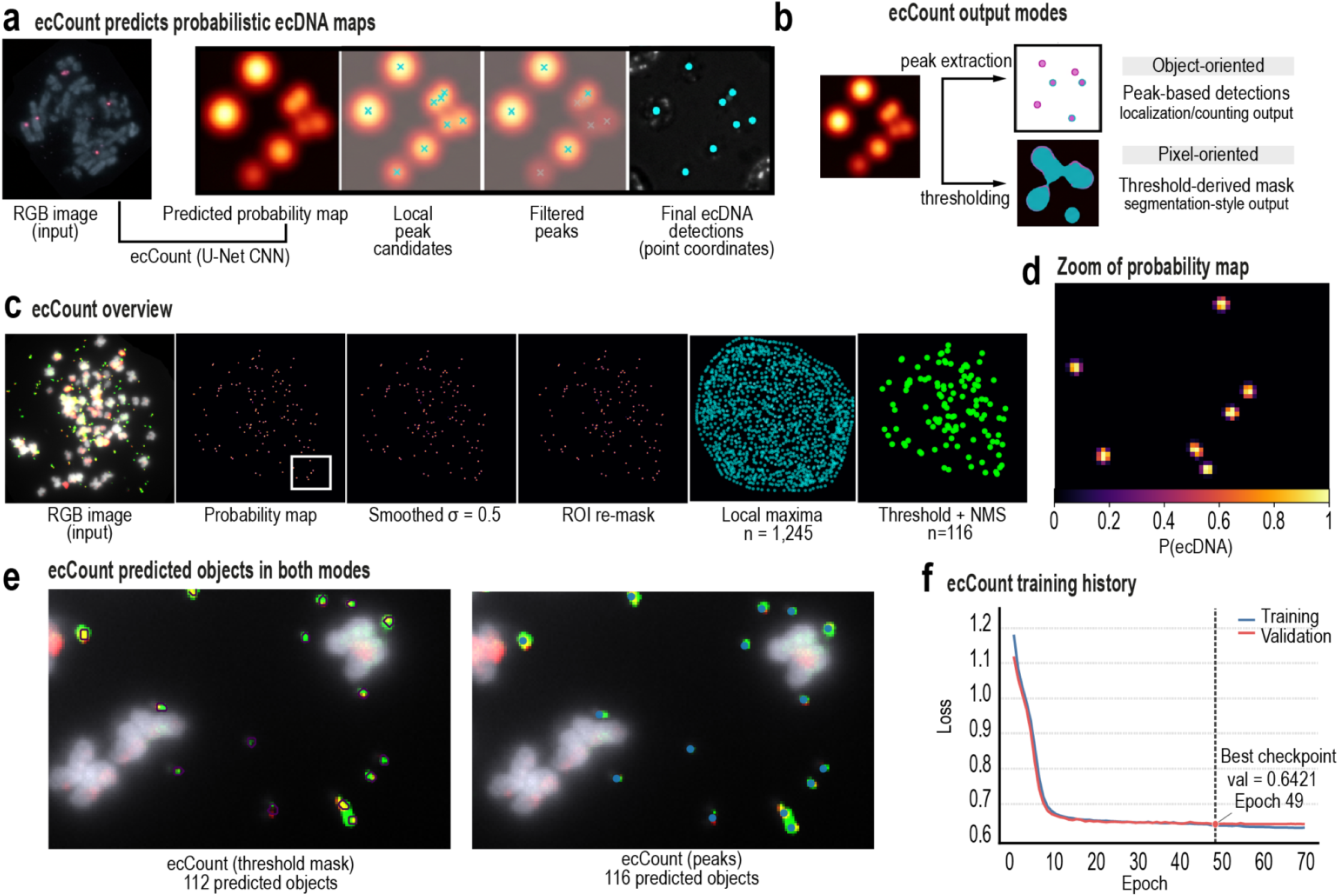
Probabilistic localization preserves individual ecDNA signals. a, ecCount reformulates ecDNA detection as probabilistic localization. An input FISH image is processed by a U-Net to generate a spatial probability map; candidate local maxima are subsequently filtered to yield ecDNA point coordinates (pixel positions of the maxima). **b,** Two modes for interpreting the ecCount probability map. Peak extraction identifies individual ecDNA locations directly for object-level localization and counting, whereas thresholding converts the same probability map into a binary, segmentation-style mask. **c,** Stepwise ecCount peak-detection workflow showing the RGB input, probability map, smoothing, ROI restriction, local-maximum detection, probability thresholding and non-maximum suppression. **d,** Enlarged probability map illustrating distinct local probability maxima corresponding to likely ecDNA positions. **e,** Comparison of the two ecCount output modes on a representative metaphase containing 116 manually annotated ecDNAs. Thresholding the probability map yields 112 connected objects, whereas peak extraction recovers 116 discrete detections. **f,** ecCount training and validation loss across 70 epochs. The final model was selected at epoch 49 on the basis of minimum validation loss before evaluation on the held-out test set.

The distinction begins with the training target. Instead of converting each manual annotation into a hard binary object with a fixed boundary, ecCount represents it as a Gaussian probability distribution centered on the annotated position^36^. Probability decreases smoothly with distance, encoding confidence in the approximate location of an ecDNA without requiring an exact object shape. Importantly, neighboring annotations can retain distinct probability maxima even when their surrounding distributions overlap, preserving closely spaced signals as separate objects.

Because ecCount predicts likely object locations rather than foreground pixels, the resulting probability map can be interpreted in two ways (**Fig. 5b**). Individual peaks can be extracted directly to generate ecDNA coordinates and counts, or the same probability map can be thresholded into a binary mask and analyzed like a segmentation output. Comparing the two readouts isolates the effect of peak extraction for a fixed trained network.

For peak-based localization, ecCount identifies local maxima from the probability map and filters them by confidence and spatial proximity (**Fig. 5c,d**). In a representative metaphase containing 116 manually annotated ecDNAs, thresholding the probability map yielded 112 connected objects, whereas peak extraction recovered 116 discrete detections (**Fig. 5e**). The final model was selected using the fixed validation partition before evaluation on held-out images (**Fig. 5f**).

By treating manual annotations as approximate locations rather than exact boundaries, probabilistic localization accommodates uncertainty in manual annotation while preserving the individual signals required for counting.

## Computational paradigms differ in accuracy and counting bias

We next compared all computational approaches on the same held-out test set to determine how differences in problem formulation translated into localization and counting accuracy (**Fig. 6a*; Extended Data Fig. 5***). Probabilistic localization performed best across the principal evaluation measures. ecCount peak detection achieved an object-level F1 of 0.939 and the lowest count error, outperforming ecCount thresholding, Label Engine, MIA, the optimized classical pipeline and ecSeg. Rankings differed across metric families: methods could place foreground pixels in approximately the correct regions yet perform poorly when those pixels failed to resolve into the correct number of discrete ecDNA objects. Pixel overlap with the annotation masks therefore did not capture the biological task of counting individual ecDNAs.

**Figure 6:**
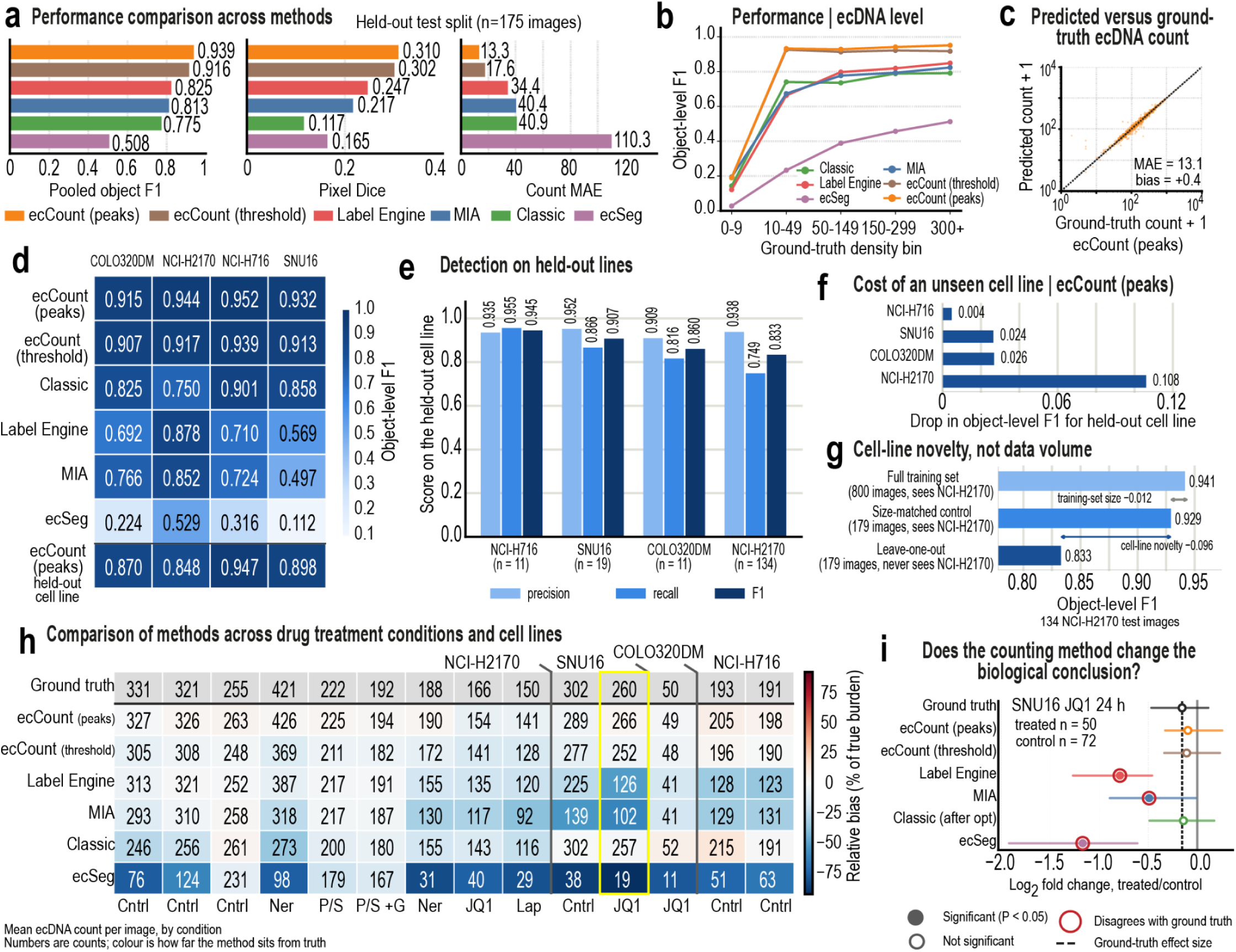
Probabilistic localization improves the accuracy, transferability and biological fidelity of ecDNA quantification. a, Comparison of all six computational outputs on the 175-image held-out test set. Performance is quantified by object-level F1 (true positives, false positives and false negatives pooled over images after one-to-one matching), pixel-level Dice and count mean absolute error. ecCount peak detection achieved the highest object-level F1 (0.939) and lowest count error (13.3 ecDNAs per image). **b**, Object-level F1 stratified by gold-standard ecDNA abundance. Performance is shown across ecDNA-count bins, revealing how detection accuracy changes with increasing ecDNA burden. ecCount peak detection maintains high performance across the major count regimes represented in the benchmark. **c**, Predicted versus gold-standard ecDNA counts for ecCount peak detection across the complete 1,145-image benchmark. The dashed line denotes perfect agreement. ecCount closely tracks gold standard across the full dynamic range, with a mean absolute error of 13.1 ecDNAs per image, a median absolute percentage error of 5% and a mean signed error (predicted minus gold-standard count, averaged over images) of +0.4. **d**, Object-level F1 across methods and cell lines, pooled over all images of each cell line in the complete 1,145-image benchmark. The appended ecCount leave-one-cell-line-out row reports each cell line scored by a model trained without it; the rows above it are scored by models developed with all four cell lines represented, and therefore include images used during training. The like-for-like comparison, in which both conditions are evaluated on held-out images, is given in **f**. **e**, Precision, recall and object-level F1 for ecCount peak detection on each cell line held out from training, scored on that cell line’s held-out test images (n = 11, 134, 11 and 19 for COLO320DM, NCI-H2170, NCI-H716 and SNU16). Precision is similar across cell lines while recall varies, indicating that the effect of an unseen cellular context is systematic under-detection rather than uniform degradation. OR matching (maximum centroid distance, 20 pixels; minimum IoU, 0.1; α = 0.5). **f**, Cost of cellular-context novelty for ecCount peak detection. For each cell line, object-level F1 for the model trained without that cell line is compared with the released model, both scored on the held-out test images of that cell line. Bars show the difference, computed at full precision. Three of the four comparisons rest on fewer than 20 images and are correspondingly imprecise. OR matching (maximum centroid distance, 20 pixels; minimum IoU, 0.1; α = 0.5). **g**, Effect of training-set size versus cellular-context novelty on ecCount performance, evaluated on the 134 NCI-H2170 held-out test images so that all three conditions are directly comparable. Reducing the training set from 800 to 179 images while retaining NCI-H2170 accounts for a smaller share of the drop than removing the cell line at the same training-set size; the two steps sum to 0.108, the NCI-H2170 value in **f**. OR matching (maximum centroid distance, 20 pixels; minimum IoU, 0.1; α = 0.5. **h**, Mean predicted ecDNA count per image for each method across drug-treatment conditions and cell lines in the complete benchmark. Values indicate mean ecDNA counts, and cell color denotes relative bias from gold standard for each condition, expressed as a percentage of ecDNA burden (blue, undercounting; red, overcounting; white, minimal bias). **i**, Effect of counting method on the inferred biological response of SNU16 cells treated with JQ1 for 24 hours. Points indicate the log2 fold change in median predicted ecDNA burden, treated versus control, with 0.5 added to each median; horizontal bars are 95% percentile bootstrap confidence intervals over 2,000 resamples. Filled markers denote P < 0.05 (two-sided Mann–Whitney U, uncorrected across contrasts) and open markers P ≥ 0.05; a red ring marks a method whose significance verdict disagrees with the gold standard. The dashed vertical line indicates the gold-standard effect size and the solid gray line indicates no change. Manual annotations show no significant change in ecDNA burden; ecCount peak detection, ecCount threshold-mask extraction and the optimized classical pipeline reproduce this conclusion, whereas Label Engine, MIA and ecSeg report a significant decrease.

All methods were less stable in images containing very few ecDNAs, where individual false-positive or false-negative calls have a large proportional effect. Across the higher-burden regimes comprising most of the benchmark, ecCount retained consistently high accuracy, whereas segmentation-based approaches varied more strongly with ecDNA burden (**Fig. 6b**). Across all 1,145 benchmark images, ecCount peak predictions closely tracked gold-standard counts over a broad dynamic range, with a mean absolute error of 13.1 ecDNAs per image (median absolute percentage error, 5%) and near-zero signed bias (**Fig. 6c*; Extended Data Fig. 6***). By contrast, segmentation-based approaches increasingly underestimated ecDNA burden as gold-standard count increased, consistent with nearby signals merging into larger foreground regions.

Performance also differed across cellular contexts (**Fig. 6d*; Extended Data Fig. 7***). NCI-H2170 contributes 888 of the 1,145 annotated metaphases, with substantially fewer examples from SNU16, NCI-H716 and COLO320DM. Several learned approaches showed marked cell-line-dependent performance, whereas ecCount maintained similar object-level F1 across all four contexts despite this imbalance (**Fig. 6d**). Leave-one-cell-line-out retraining, performed for ecCount only, showed transfer to cell lines absent from training: on their held-out test images, object-level F1 was 0.004–0.108 lower than that of the released model (**Fig. 6e–g*; Extended Data Fig. 8***). Within the benchmark, probabilistic localization was the paradigm least sensitive to ecDNA burden and cellular context (**Fig. 6b,d**).

## Detection bias alters biological interpretation of ecDNA dynamics

A central application of automated ecDNA imaging is to quantify how ecDNA populations change under perturbation. Drug treatment can substantially reshape ecDNA abundance, and distinguishing true shifts in copy number from measurement error is essential for interpreting these changes as loss, selection or adaptation. We therefore compared computational and manual counts across multiple cell lines and perturbations, including antibiotic agents, tyrosine kinase inhibitors (lapatinib, neratinib) and chromatin-modifying agents (JQ1). Counting bias differed substantially among methods and experimental conditions, whereas ecCount peak detection most closely preserved gold-standard ecDNA burden (**Fig. 6h**).

These differences were sufficient to alter biological inference. In SNU16 cells treated with JQ1 for 24 hours, manual annotations showed no significant change in ecDNA abundance. ecCount and the optimized classical pipeline reproduced this result, whereas Label Engine, MIA and ecSeg each reported a significant decrease (**Fig. 6i**). Analysis restricted to held-out images reproduced the broader pattern across treatment conditions and cell lines (Extended Data Fig. 9a–c). Thus, count-dependent error can become experimentally indistinguishable from a change in ecDNA biology, making quantitative fidelity essential when automated imaging is used to measure ecDNA dynamics.

## Discussion

The ability to measure ecDNA has not kept pace with the rapidly expanding biology of ecDNA. Existing computational approaches have demonstrated that automation is possible, but the field lacks an open reference imaging resource on which methods can be trained, tested and improved. This represents an important bottleneck for a field that has rapidly emerged as a major challenge in cancer biology: we increasingly understand that ecDNA can shape tumor evolution, heterogeneity and therapeutic resistance, but our ability to measure these dynamics directly across large cell populations remains limited.

The rapid development of computer vision provides an opportunity to remove this bottleneck^24,33,34,37^. Previous methods^4,19,21^ have treated metaphase FISH images largely through rule-based image processing or semantic segmentation. In the absence of shared images, annotations and evaluation standards, however, it has been difficult to determine how these approaches compare, where they fail or whether advances in computer vision can improve upon them. We therefore built a reference resource that brings together nearly 3,000 native-resolution metaphase FISH images with annotations, benchmark partitions, model predictions and openly extensible computational workflows. Using this common resource, we compared across computational paradigms on the same underlying imaging problem. This comparison revealed that the most consequential difference between approaches was not simply accuracy, but measurement bias.

Rule-based and segmentation-based approaches systematically underestimated ecDNA abundance, and the magnitude of undercounting increased with ecDNA burden. This is more consequential than a constant counting error because cells and experimental conditions with different ecDNA burdens are distorted by different amounts. High-copy cells are disproportionately undercounted, compressing the measured range of ecDNA abundance and reducing apparent population heterogeneity. When ecDNA burden changes between conditions, this count-dependent bias can exaggerate, diminish or even create an apparent biological difference. This consequence was evident in the perturbation experiments, where segmentation-based counting changed the inferred treatment response in cases for which the manual annotations showed no significant change in ecDNA burden. For quantitative ecDNA studies, the relevant standard is therefore not simply whether an algorithm detects ecDNA accurately, but whether it preserves differences in ecDNA burden across cells and experimental conditions.

Probabilistic localization substantially reduced this bias by changing the way the problem is posed. Rather than asking a network to reproduce the boundaries of ecDNA objects from imperfect manual annotations, ecCount learns where individual signals are likely to occur and resolves them as discrete probability peaks. It reflects a closer alignment between the computational task and the biological measurement: the number and position of individual ecDNA signals.

The resulting capability changes what metaphase FISH can contribute to ecDNA biology. Automated measurements that remain quantitatively stable across ecDNA burden create a path toward following copy-number distributions through drug treatment, withdrawal and adaptation; identifying rare cellular states that persist under selective pressure; and determining how rapidly populations reconstruct ecDNA heterogeneity. The same framework can expand to multicolor FISH, where individual ecDNA species and their co-occurrence can be measured, and ultimately to other imaging contexts. Importantly, the resource makes these advances extensible: new images, annotations and algorithms can be added to a common benchmark rather than developed against inaccessible training sets and incomparable performance measures. The present resource does not resolve every challenge. Its benchmark derives from one laboratory and imaging platform. Leave-one-cell-line-out analyses show that ecCount transfers to previously unseen cell lines, but performance improves when annotated images of the target line are included: on NCI-H2170 test images, a 179-image training set containing the line reached an object-level F1 of 0.929, against 0.833 for an equally sized set without it and 0.941 for all 800 training images (**Fig. 6g**). Generalization across laboratories, microscopes and probe chemistries, however, remains to be established.

Manual annotation remains an imperfect reference, and future contributions from additional laboratories, microscopes and probe chemistries will be important for defining the limits of transferability. These limitations also underscore the purpose of an open resource. Rather than treating a single model as a finished solution, the images, annotations and evaluation framework provide a foundation on which the community can test new methods, expose failure modes and progressively broaden the range of images that can be quantified reliably. To facilitate this expansion, we also provide a series of interactive Jupyter notebook tutorials that guide users through preparing their own annotated images, retraining the open-source models and evaluating their performance within the same benchmarking framework. In this way, the resource is designed not only to make the current models reproducible, but to allow them to be adapted, challenged and improved as new imaging datasets become available. More broadly, scaling ecDNA imaging requires a shift from automating what a human sees to preserving what an experiment measures. For ecDNA, that measurement is the distribution of discrete DNA copies across individual cells. By making the underlying imaging data accessible, establishing a common standard for evaluating computational approaches and reducing count-dependent measurement bias, this work provides the infrastructure needed to move metaphase FISH from labor-intensive enumeration toward scalable quantitative analysis of ecDNA dynamics.

## Data availability

All images, annotations and benchmark materials generated in this study are available from the BioImage Archive under accession S-BIAD4097 (https://doi.org/10.6019/S-BIAD4097), released under CC BY 4.0. The deposition comprises 2,986 paired probe-channel RGB and DAPI image sets; the manual ecDNA annotations (gold standard; the archive’s “ground truth” section) as point coordinates, as binary masks in which each point is rendered as a 5 × 5-pixel diamond, and as the same masks in sparse NumPy (.npz) format; manual region-of-interest masks for the 1,145-image benchmark subset; model-predicted region-of-interest masks for all 2,986 images; prediction masks from every compared method; metadata tables; and the fixed training, validation and held-out test partitions. Each image set carries a unique identifier linking every modality and annotation.

## Code availability

Source code for the classical computer-vision pipeline, ecCount, region-of-interest prediction and the unified evaluation framework is available at https://github.com/Brunk-Lab/ecdna-bench under the MIT License; Label Engine is available at https://github.com/Brunk-Lab/Label-Engine. The repository contains model-training and inference workflows, fixed configuration files, benchmark-evaluation scripts, the adapters that convert each comparator’s native output (ecSeg label maps, MIA and Label Engine masks) to the common binary-mask format, Jupyter notebook tutorials and the scripts that generate every figure’s source data. Trained model weights and frozen benchmark predictions are distributed as release assets.

## Methods

### Primary cell culture

NCI-H2170, SNU16, COLO320DM and NCI-H716 cells were obtained from ATCC and cultured in RPMI 1640 medium (Gibco) supplemented with 10% heat-inactivated fetal bovine serum (Gibco). SUM159PT cells were obtained from the laboratory of Gary Johnson (University of North Carolina at Chapel Hill) and cultured in Ham’s F-12 medium supplemented with 5% fetal bovine serum, 10 mM HEPES, 1 μg ml⁻¹ hydrocortisone and 5 μg ml⁻¹ insulin. Cells were maintained at 37 °C in a humidified incubator with 5% CO₂ and used within three passages after thawing. SUM159PT cells were authenticated by short-tandem-repeat profiling and tested for mycoplasma contamination before use. Adherent cells were harvested with 0.25% trypsin in DPBS (Gibco). Viable cells were stained with trypan blue (Invitrogen, T10282) and counted using a Countess 3 automated cell counter (Invitrogen).

### Metaphase sample preparation and imaging

Cells at approximately 70% confluency were arrested in metaphase by treatment with 0.1 μg ml⁻¹ colcemid (FUJIFILM Irvine Scientific) for 12–20 hours. Adherent cells were detached by trypsinization, collected and centrifuged before incubation in 0.075 M KCl (Gibco) for 15 min at 37 °C to induce hypotonic swelling. Cells were fixed by dropwise addition of freshly prepared modified Carnoy’s fixative (3:1 methanol acetic acid). Fixation was repeated three times, and the final cell suspension was adjusted to approximately 6 × 10⁶ cells ml−1.

Metaphase slides were prepared on Superfrost microscope slides (Fisherbrand, 12-550-123) by dropping the fixed cell suspension onto the slide and allowing it to air dry for 1 hour. Slides were equilibrated in 2× saline-sodium citrate (SSC; Invitrogen, 15557-036) and dehydrated through 70%, 85% and 100% ethanol.

DNA FISH was performed using locus-specific probes obtained from Empire Genomics. Probes targeting amplified loci including ERBB2 (17q12), MYC (8q24), CDC6 (17q21.2) and PVT1 (8q24) were conjugated to spectrally distinct fluorophores as appropriate for each cell line and experiment. Hybridization mixture (5 μl) was applied to each slide beneath a coverslip, and slides were denatured at 72 °C for 2 min and hybridized for 16–20 hours at 37 °C in a humidified chamber. Slides were subsequently washed in 0.4× SSC followed by 2× SSC containing 0.05% Tween-20 and mounted using SlowFade Diamond Antifade Mountant with DAPI (Invitrogen, S36964).

### Fluorescence imaging

Metaphase spreads were imaged using an Echo Revolution automated fluorescence microscope at ×60 magnification. Probe-channel and DAPI images were acquired for each metaphase at a native resolution of 2,448 × 2,048 pixels. Images were acquired from the same slide or replicate slides prepared from the same metaphase preparation to minimize experimental variation in imaging conditions. Images were visualized and manually inspected using Fiji/ImageJ (v.2.1.0/1.53c)^38^.

### Imaging-resource assembly and quality control

Metaphase FISH images were curated into a reference resource by assigning each metaphase spread a unique identifier and matching this identifier across probe-channel RGB images, DAPI images and annotation files. The final resource contains 2,986 complete image sets from five ecDNA-positive cancer cell lines^6,25–27^: NCI-H2170 (n = 1,891), SUM159PT (n = 458), SNU16 (n = 273), COLO320DM (n = 225) and NCI-H716 (n = 139). Each image set contains matched RGB and DAPI images together with ecDNA annotation files.

Automated quality-control procedures were used to confirm unique identifiers, complete modality coverage and consistent spatial dimensions across the resource. Images were not excluded on the basis of ecDNA abundance, signal-to-noise ratio or image complexity, allowing the released resource to retain the heterogeneity encountered during routine metaphase FISH analysis.

### Manual ecDNA annotation

ecDNA signals were manually annotated on paired probe-channel RGB and DAPI images with the point tool of Fiji/ImageJ^38^. Annotators used probe-specific fluorescence to identify amplified loci and the DAPI signal to determine chromosomal context, and placed one point at the approximate center of each fluorescent signal judged to represent extrachromosomal amplification. No minimum ecDNA size threshold was imposed. Annotations were made by multiple trained operators without a consensus procedure or confidence scores, so they retain the spatial and interpretive uncertainty of manually classifying small or crowded FISH signals. Inter-annotator variability under the same protocol and imaging platform was quantified in the MIA study^19^ and was not re-measured here. We refer to these manual annotations as the gold standard.

Annotations are released in three formats: the point coordinates; binary masks in which each point is rendered as a 5 × 5-pixel diamond (13 pixels) centered on the annotation; and the same masks as sparse NumPy (.npz) arrays. Touching diamonds form a single connected component, and the 8-connected components of the rendered mask (minimum area 3 pixels) define the gold-standard objects and counts used throughout (***Extended Data Fig. 1d***). The diamonds encode position, not object extent; they served as training targets for the segmentation models and as the reference for pixel-level metrics.

### Manual metaphase region-of-interest annotation

Metaphase FISH fields can contain unburst nuclei, neighboring cells, debris and other features unrelated to the metaphase being quantified. Each benchmark image therefore has a manually drawn region of interest (ROI): a single freehand region enclosing the metaphase spread to be quantified, drawn with a lasso tool to enclose all annotated ecDNA of the spread while excluding unannotated material. Pixels outside the ROI were set to zero for every method, and all benchmark predictions and evaluations were restricted to it (**Fig. 2c**). Manual ROIs exist for 1,145 of the 2,986 image sets; the other 1,841, including all SUM159PT images, carry point annotations but no manual ROI and are released with predicted ROIs (see *Automated ROI prediction*), outside the benchmark.

### Benchmark construction and dataset partitions

The ROI-constrained benchmark contains 1,145 manually annotated images from four cell lines: NCI-H2170 (n = 888), SNU16 (n = 123), NCI-H716 (n = 70) and COLO320DM (n = 64). SUM159PT was retained in the complete resource but excluded from the primary benchmark because manual ROI masks were not available.

Benchmark images were partitioned at the unique-image-identifier (metaphase) level into fixed training (n = 800), validation (n = 170) and held-out test (n = 175) sets, stratified by cell line in approximately 70:15:15 proportions (Supplementary Table 1). All image modalities, ROI masks and annotations belonging to a metaphase were assigned to the same partition, and pairwise overlap checks confirmed that the partitions were mutually exclusive. Images from the same slide or treatment condition can fall into different partitions, so the test partition measures generalization within the imaged conditions; transfer to an unseen cellular context was assessed separately (*Leave-one-cell-line-out generalization*). The partitions were used by every model fitted in this study (the classical pipeline, Label Engine and ecCount); ecSeg and MIA were not retrained (see *ecSeg and MIA comparator models*).

Gold-standard ecDNA burden in the benchmark ranged from 1 to 1,687 objects per image, with a mean of 199.2 and median of 159. For stratified analyses, images were grouped by gold-standard count into bins of 0–9, 10–49, 50–149, 150–299 and ≥300 ecDNA objects per image (count bins).

### Classical computer-vision pipeline

A rule-based computer-vision pipeline with 10–13 tuned parameters per cell line was developed as a baseline for ecDNA detection. Analysis was restricted to the manually defined metaphase ROI. Probe-channel RGB images were collapsed into a single grayscale image using a channel-wise maximum-intensity transformation. Images were then processed by an ordered sequence of three or four operators, selected automatically as described below, from background suppression, top-hat transformation^39^, gamma correction, unsharp masking, bilateral filtering^40^, contrast-limited adaptive histogram equalization (CLAHE)^41^ and sigmoid intensity transformation.

Candidate foreground objects were identified by Otsu thresholding^42^ scaled by a tuned factor k, followed by morphological closing and eight-connected-component extraction^30^. Components outside a tuned area range were discarded, and components whose centroids lay within a tuned merge distance were merged. Each remaining object was then classified from its mean color in hue–saturation–value (HSV) space: objects with mean value above, and mean saturation below, cell-line-specific thresholds — bright, desaturated, chromosome-associated signal — were labeled chromosome-like and discarded, and all others were counted as ecDNA.

Pipeline parameters were optimized independently for each cell line in three stages, each scored by pooled object-level F1 under the canonical matching (see Object-level matching). Stage 1 exhaustively evaluated all 1,050 ordered sequences of three or four of the seven operators, with default operator parameters, on 25 training images per cell line, and kept the sequence with the highest F1. In stage 2, Gaussian-process Bayesian optimization^31^ was used to optimize the parameters of the selected operators together with the two HSV thresholds. Stage 3 then optimized the Otsu factor k, the closing-kernel size, the minimum and maximum object area and the merge distance, with preprocessing held fixed. Stages 2 and 3 used a Gaussian-process surrogate with an upper-confidence-bound acquisition (bayesian-optimization v3.1.0; Matérn-5/2 kernel, κ = 2.576) and pooled F1 on up to 100 training images per cell line as the objective, and the evaluated configuration with the highest validation F1 was retained. Test images were used for neither optimization nor selection; their F1 was recorded for reporting only. Final detection parameters are provided in Supplementary Table 3; the complete parameter-search space and cell-line-specific preprocessing operator sequences are provided in default_classical_optimised.yaml, released with the code.

### Label Engine segmentation model

Label Engine is an open-source, retrainable segmentation framework for ecDNA detection, developed here and used in its automatic mode, which takes the image alone: no prompts were supplied during training or inference. The network is a U-Net^32^ with a residual convolutional image encoder and a residual decoder. The encoder processes the native-resolution RGB FISH image and produces a five-scale feature pyramid. The decoder restores spatial resolution by transposed convolution and combines each upsampled level with the corresponding encoder features through skip connections, and a final prediction head outputs a two-channel softmax map corresponding to background and ecDNA.

The encoder contains an initial convolutional block followed by four downsampling stages, with feature widths of 16, 32, 64, 128 and 256 channels; the deepest representation has a stride of 16 relative to the input image. Training used native-resolution RGB images from the benchmark training partition. Data augmentation included random flipping, geometric transformations, brightness and contrast perturbation and channel-wise intensity jitter, applied synchronously across model inputs.

Label Engine was optimized using Adam^43^ with an initial learning rate of 5 × 10⁻⁵ and a MultiStep learning-rate schedule with milestones at iterations 1,000 and 1,500. Training was performed for 400 epochs using focal loss^44^ with equal class weighting and deterministic initialization. At inference the network received the image alone, and the ecDNA probability channel was thresholded at 0.4 to give the binary mask used in the common evaluation framework. Label Engine also provides an interactive mode in which user clicks guide the prediction; that mode is under development and was not used in this study.

### ecSeg and MIA comparator models

ecSeg^21^ and ecDNA-MIA^19^ were included as previously developed comparators; neither was retrained on the benchmark partitions. ecSeg is distributed as a trained Keras/TensorFlow network (metaseg.h5; architecture and weights) with inference code, but without training code or training data. The network classifies each pixel of the DAPI channel as background, nucleus, chromosome or ecDNA, and its post-processing uses the nucleus and chromosome classes. Our annotations mark only probe-defined ecDNA and therefore cannot supervise this four-class output: retraining would require pixel labels for nuclei and chromosomes, which the resource does not contain, or a new output head, which would no longer be the published model. ecSeg was therefore applied zero-shot with its released weights, and its results measure transfer of the published model to new imaging data rather than the capacity of its architecture. For MIA, our previously developed deep-learning framework, we used archived prediction masks generated using the top-performing configuration established in the original study.^19^

ecSeg was evaluated using the pretrained metaseg.h5 model^21^ on the ROI-masked RGB images; its preprocessing uses the third (blue) channel, which carries the DAPI signal in these images, and its segmentation step has no user-set parameters. Predicted class-index maps were converted to binary ecDNA masks by selecting class index 3, which corresponds to ecDNA in the metaseg.h5 label scheme. This assignment was independently verified by comparing the number of ecDNA objects recovered from the class-3 masks with ecSeg’s internal quantification output, with perfect agreement across all images (mean absolute error = 0).

MIA prediction masks were generated previously using the top-performing configuration established in the original study and were not regenerated or retrained for the present benchmark. Predictions were available as single-channel PNG images encoded as {0,1} or {0,255}; no additional probability threshold was applied.

MIA and ecSeg predictions were processed through the same harmonization framework before evaluation. Any non-zero pixel was treated as foreground, connected components smaller than 3 pixels were removed, and binary masks were written at the original image resolution. Neither method received method-specific post-processing. The MIA harmonization workflow is implemented in ecdna_bench.baselines.mia in the released software package. All comparator outputs were subsequently stored in a common format for evaluation. The previously published ECdetect method^4^ was not included as an executable comparator because runnable code was not available.

### ecCount probabilistic-localization model

ecCount was developed to treat manual ecDNA annotations as uncertain object locations rather than exact object boundaries. ROI-masked RGB images were used as model inputs, with pixels outside the manually defined metaphase ROI set to zero. Images were downsampled by a factor of two in each dimension, to 1,024 × 1,224 pixels (height × width; area interpolation), and 8-bit intensities were divided by 255 without per-image normalization; predicted coordinates were scaled back by two to the native frame for evaluation. Spatial transformations were applied synchronously to images, annotation targets and ROI masks.

### Gaussian localization targets

Each annotated ecDNA position was represented by an isotropic Gaussian target centered on the centroid of each gold-standard object,

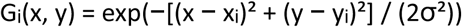

where (xᵢ, yᵢ) denotes the annotated ecDNA location. The final training target was defined as the element-wise maximum across individual Gaussian peaks and normalized to [0, 1]. Taking the maximum rather than the sum keeps every annotated position at the same target height regardless of how many neighbors it has; summing overlapping Gaussians, as in density-estimation approaches that recover counts by integrating the map^36^, would instead raise the target where annotations are dense, so intensity would encode local crowding rather than position. Because ecCount recovers objects by locating maxima rather than by integrating intensity, the maximum also preserves one peak per annotation where neighboring signals overlap. Gaussian width was optimized using the validation set, with σ = 1.0 pixel at model resolution (two native pixels) selected for the final model.

### Network architecture and training

ecCount uses a U-Net implemented in PyTorch^45^. The model accepts a three-channel ROI-masked RGB image and outputs a single-channel logit map at the same spatial resolution. The encoder contains an initial double-convolution block followed by four downsampling stages, and the decoder contains four upsampling stages with skip connections and a final 1 × 1 convolution. Convolutional blocks use 3 × 3 convolutions, GroupNorm^46^ and ReLU activation. The final architecture uses 32 base channels, eight normalization groups, bilinear upsampling, no dropout and 7,849,601 trainable parameters.

Training minimized a composite loss combining weighted binary cross-entropy and soft Dice loss^47^. The task is heatmap regression rather than segmentation: the cross-entropy term is computed on the logits against the graded Gaussian target, with a positive-class weight that offsets the foreground–background imbalance, and the Dice term acts on the same graded target. The Dice term was retained because the equally weighted combination outperformed cross-entropy-only and Dice-only training, each at positive-class weights of 10, 20 and 50, on pooled validation F1 (Supplementary Methods §10.6); mean-squared-error and focal-type heatmap losses were not evaluated. With positive-class weight w, the per-pixel minimizer of the cross-entropy term is wt/(1 − t + wt) for target value t, a monotone transform of the target, so the output preserves the positions of target maxima but is not a calibrated probability. The final configuration used a positive-class weight of 20, equal weighting of the binary-cross-entropy and Dice terms and smoothing of 1 × 10−6. Optimization used Adam^43^ with an initial learning rate of 1 × 10−4, batch size 2 and no weight decay. Augmentation included horizontal and vertical flipping and multiplicative brightness jitter. The final model was trained for 70 epochs using ReduceLROnPlateau with patience 5 and factor 0.5. The checkpoint with minimum validation loss was selected at epoch 49 before held-out evaluation.

### Peak-based inference

At inference, the network’s logit map was passed through a sigmoid to give the probability map, with values in [0, 1], which was smoothed with a Gaussian kernel and re-restricted to the metaphase ROI. Candidate ecDNA locations were identified as local probability maxima^30^ and filtered by an absolute probability threshold followed by greedy Euclidean non-maximum suppression^48^. The frozen final parameters were threshold_abs = 0.35, peak_min_distance = 2, nms_min_distance = 2, smooth_sigma = 0.5 and exclude_border = 0.

Two outputs were generated from the same ecCount probability map. For peak-based localization, retained local maxima were exported as integer pixel coordinates, scaled to the native frame and rendered as 5 × 5-pixel diamonds, the footprint of the gold-standard points, for object extraction. For segmentation-style comparison, the probability map was thresholded at 0.5 to produce a binary mask. These outputs were evaluated independently as ecCount peaks and ecCount threshold, allowing the effect of output interpretation to be compared while holding the trained network and underlying probability map constant.

### ecCount hyperparameter optimization

ecCount hyperparameters were selected using the validation partition without access to the held-out test set. Optimization proceeded in stages. Post-processing parameters were first evaluated across 108 candidate configurations. Gaussian target width was then evaluated for σ values of 0.5–3.0 pixels using independently trained models. Loss weighting was evaluated across positive-class weights of 10, 20 and 50 and alternative binary-cross-entropy and Dice-loss weights. Finally, training duration and learning-rate scheduling were evaluated across 30-, 50- and 70-epoch schedules. The final configuration was frozen before evaluation on the held-out test partition.

### Leave-one-cell-line-out generalization

To measure how ecCount behaves on a cell line absent from its training data, the model was retrained from scratch four times, once per benchmark cell line. In each run, every image of one cell line was removed from both the fixed training and validation partitions, and the trained model was then scored on all images of that cell line. Training sets contained 756 images (COLO320DM held out), 179 (NCI-H2170), 751 (NCI-H716) and 714 (SNU16), with corresponding validation sets of 161, 37, 160 and 152 images. Evaluation sets were the 64, 888, 70 and 123 images of the held-out cell line, which together reconstitute the 1,145-image benchmark and its 228,039 annotated ecDNA objects. Split composition was verified against the frozen benchmark manifest before training, and no image contributed to both model development and evaluation within a run.

Removing NCI-H2170 removes 78% of the training data as well as the cell line, confounding cellular-context novelty with training-set size. A fifth run was therefore added as a size-matched control. Its training and validation partitions were drawn by stratified sampling across all four cell lines to the same sizes as the NCI-H2170 leave-one-cell-line-out run (179 training and 37 validation images), so that the two runs differ in composition but not in volume. Because the control was exposed to NCI-H2170 training and validation images, it was scored on the 134 NCI-H2170 held-out test images only; for the comparison in Fig. 6g, the NCI-H2170 leave-one-cell-line-out run was rescored on the same 134 images.

All five runs used the architecture, optimization settings and schedule of the released ecCount model: 7,849,601 trainable parameters, inputs resampled to 1,024 × 1,224 pixels, batch size 2, Adam with an initial learning rate of 1 × 10⁻⁴ and ReduceLROnPlateau scheduling (patience 5, factor 0.5), 70 epochs, single precision and a fixed random seed. The retained checkpoint was selected by minimum validation loss within each run. Because validation partitions differ between runs, validation losses index checkpoint selection within a run and are not comparable across runs. Peak-based inference used the frozen parameters of the released model (threshold_abs = 0.35, peak_min_distance = 2, nms_min_distance = 2, smooth_sigma = 0.5, exclude_border = 0), and all scoring used the canonical OR matching policy (maximum centroid distance, 20 pixels; minimum IoU, 0.1; α = 0.5).

Two quantities are reported for each cell line, both evaluated on the same images. The unseen score is the leave-one-cell-line-out model evaluated on the held-out test images of the excluded cell line (n = 11, 134, 11 and 19 for COLO320DM, NCI-H2170, NCI-H716 and SNU16). The in-distribution score is the released model evaluated on those same images, which it likewise did not train on. Restricting both to a common evaluation set leaves the presence or absence of the cell line during training as the only difference between them. The cost of cellular-context novelty reported in Fig. 6f and Extended Data Fig. 8e is the difference between these two quantities, computed at full numerical precision before rounding for display. Leave-one-cell-line-out models were additionally scored on every image of the excluded cell line (n = 64, 888, 70 and 123); those values appear in the appended row of Fig. 6d, which pools over all images of each cell line for every method. Three of the four common evaluation sets contain fewer than 20 images, so the corresponding differences are correspondingly imprecise.

Each configuration was trained once at a fixed seed. Replicate runs were not performed, so differences of comparable magnitude between cell lines are not resolved by these experiments.

### Automated ROI prediction

To extend the resource beyond manually annotated metaphases, we trained an auxiliary ROI-prediction model using the 1,145 manually defined ROI masks. The model is a four-channel residual U-Net^32,49^ with 35,923,337 parameters and was trained using the same 800-image training and 170-image validation partitions used for ecCount; the 175-image held-out test set was not used during model development. Architecture, optimization and post-processing are described in Supplementary Methods §11.

The trained model was applied to all 2,986 images in the resource, including the 1,145-image benchmark subset, for which both manual and predicted ROI masks are provided. Across the benchmark, predicted ROIs showed a median intersection over union of 0.821 and median Dice coefficient of 0.902 relative to manual regions, while retaining 98.5% of manually annotated ecDNA objects. Regions omitted by the predicted masks contained substantially fewer ecDNA annotations than retained regions, indicating that most geometric disagreement occurred in areas with little ecDNA signal. Agreement between predicted and manual regions is reported in Supplementary Methods §11.8.

Predicted ROIs were not used for primary model training, benchmarking or performance comparisons in this study; these analyses used manually defined ROI masks exclusively. Predicted masks and the ROI-prediction workflow are released as separately identified computational annotations to support application of the resource and future fully automated ecDNA analysis. As an end-to-end test, the released ecCount model was also run inside the predicted ROIs of the benchmark images and scored against all annotations (Supplementary Methods §11.13).

### Harmonization of model outputs

The evaluated approaches produce object lists with bounding boxes, binary masks or point coordinates. Every output was therefore converted to a binary mask at the native 2,448 × 2,048-pixel resolution before comparison. Point outputs were rendered as 5 × 5-pixel diamonds, and all masks were decomposed into 8-connected components^30^ of at least 3 pixels; each component is one object with a centroid, used for the distance criterion, and a pixel footprint, used for the overlap criterion. Gold-standard objects were the components of the rendered annotation masks, which were also used for the pixel-level metrics.

### Object-level matching

Predicted and gold-standard objects were matched one-to-one in two steps: an eligibility rule that decides which pairs may be matched, and an assignment step that chooses among the eligible pairs. For each pair, the centroid distance and the mask intersection over union (IoU) were computed, and eligible pairs were assigned one-to-one with the Hungarian algorithm^50^.

A pair was eligible if its centroid distance was at most d_max = 20 pixels or its mask IoU was at least 0.1 (OR policy); the stricter AND policy required both. Eligible pairs were then assigned at minimum total cost C_ij = α(1 − IoU_ij) + (1 − α)·d_ij/d_max with α = 0.5, so the weighting ranks competing eligible pairs but does not decide eligibility. In practice, eligibility under OR was set by the distance criterion: at d_max = 20 pixels, object-level F1 on the held-out test set was identical for every minimum IoU from 0.05 to 0.5. We used OR because the gold-standard objects are fixed 5 × 5-pixel diamonds rather than object outlines, so an overlap requirement penalizes correctly placed detections whose rendered shape differs; under AND, F1 was governed by the minimum IoU (***Extended Data Fig. 6e***).

Matched pairs were classified as true positives, unmatched gold-standard objects as false negatives and predictions without an eligible gold-standard partner as false positives. Additional predictions that were eligible for the same gold-standard object but were not selected during one-to-one assignment were tracked separately as duplicate detections and were not counted as additional false positives.

To assess sensitivity to the matching definition, we repeated evaluation across centroid-distance thresholds of 5–100 pixels and IoU thresholds of 0–0.5 under both OR and AND policies, where AND required both criteria to be satisfied. Absolute F1 changed across matching conditions. Under OR, the two ecCount outputs ranked first and second at all 70 grid points on the held-out test set, whereas the order of the other methods varied at some grid points; under AND, the ranking depended on the minimum IoU. Full sensitivity analyses are provided in Extended Data Fig. 5g and Extended Data Fig. 6a,e, and in Supplementary Methods §13.

### Performance metrics and statistical analysis

Performance was evaluated at object, pixel and count levels. Object-level precision, recall and F1 = 2TP/(2TP + FP + FN) were computed from matched (TP), unmatched predicted (FP) and unmatched gold-standard (FN) objects, so object-level F1 is the Dice coefficient computed over objects. Pixel-level precision, recall, intersection over union and Dice coefficient (hereafter Dice) were computed between the harmonized binary masks and the rendered gold-standard masks; because the latter are 5 × 5-pixel diamonds, Dice measures agreement with this rendering rather than with object outlines and is a secondary metric. Count metrics were the mean absolute error, root-mean-square error, mean signed error (bias; predicted minus gold-standard count, averaged over images), Pearson and Spearman correlations and the median absolute percentage error for images with non-zero gold-standard counts.

Object-level and pixel-level true positives, false positives and false negatives were pooled across images before the metrics were computed (micro-averaging). Count metrics were computed per image and averaged. Performance was additionally stratified by cell line and by gold-standard count bin.

Primary comparative performance claims were based on the 175-image held-out test set. Analyses explicitly described as using the full benchmark included all 1,145 ROI-annotated images.

Paired model comparisons were performed using two-sided Wilcoxon signed-rank tests on per-image object F1, pixel-level Dice and absolute count error. Simultaneous comparisons across models used the Friedman test, and multiple testing across these paired comparisons was controlled with the Benjamini–Hochberg procedure at a false-discovery rate of 5%. Treatment effects were quantified as the log2 ratio of the median predicted ecDNA count in treated cells to that in matched control cells, with 0.5 added to each median so that the ratio remains defined when a method predicts a median of zero. Ninety-five percent confidence intervals were obtained by percentile bootstrap over 2,000 resamples, each arm resampled independently with replacement to its own size, using numpy.random.default_rng reseeded to 0 for every contrast. Treated-versus-control differences were tested with two-sided Mann–Whitney U tests and called significant at P < 0.05; no multiple-comparison correction was applied across contrasts. A contrast was evaluated only where both arms contained at least eight images across the full benchmark, or at least five images within the held-out validation and test split. Statistical analyses were performed on predictions generated from frozen models and parameter configurations; no model tuning was performed using the held-out test set.

### Software environment and reproducibility

Computational analyses were performed on a high-performance computing cluster. Separate Conda environments were maintained for the classical computer-vision pipeline, Label Engine, ecSeg, MIA and ecCount to preserve model-specific software dependencies.

Random seeds and fixed dataset partitions were used where applicable, and intermediate outputs were retained to support reproducibility. These outputs include trained model checkpoints, probability maps, harmonized predictions, predicted object coordinates, per-image performance metrics, benchmark summaries and figure source data. Model parameters and evaluation settings were frozen before held-out test evaluation.

### Tutorials

To facilitate reuse, we provide interactive Jupyter notebook tutorials that guide users through image and annotation preparation, model training and retraining, inference and performance evaluation, so that the released workflows can be applied to new FISH images and new models can be evaluated in the same benchmark framework.

## Acknowledgements

The authors thank the UNC Flow Cytometry Core, with special acknowledgment to Janet Dow and Ramiro Diz, for their expertise and support. We also extend our gratitude to staff of the UNC Microscopy Core for their valuable contributions. P.B. was supported by funding from the PhRMA Foundation and the Lung Cancer Research Foundation Boehringer Ingelheim Early Career Investigator Award. E.B. was supported by funding from the PhRMA Foundation and the Lung Cancer Research Foundation Boehringer Ingelheim Early Career Investigator Award. M.M. was supported by funding from the Lung Cancer Research Foundation Boehringer Ingelheim Early Career Investigator Award.

## Author contributions

Conceptualization, E.B. ; methodology, P.B., R.S., Q.L., M.N., E.B. ; formal analysis, P.B., R.S., Q.L., A.I., E.B.; funding acquisition, E.B.; investigation, P.B., R.S., Q.L., E.B.; resources, E.B., M.N. ; supervision, E.B., M.N. ; validation, P.B., R.S. ; visualization, P.B., E.B.; writing—original draft, P.B., E.B.; writing—review and editing, All authors. All authors have read and agreed to the published version of the manuscript.

## Inclusion and Ethics Statement

All experiments involving cell lines were conducted in compliance with institutional biosafety and research integrity guidelines approved by the University of North Carolina at Chapel Hill. No human or animal subjects were involved in this study. We are committed to fostering a supportive research environment. Our team reflects a range of backgrounds and training levels, and we actively mentor junior scientists across disciplines. We strive to ensure that our research practices are transparent, reproducible, and accessible to the broader scientific community. Data and code from this work are openly shared to promote collaboration and accountability.

## Extended Data Figures

**Extended Data Figure 1:**
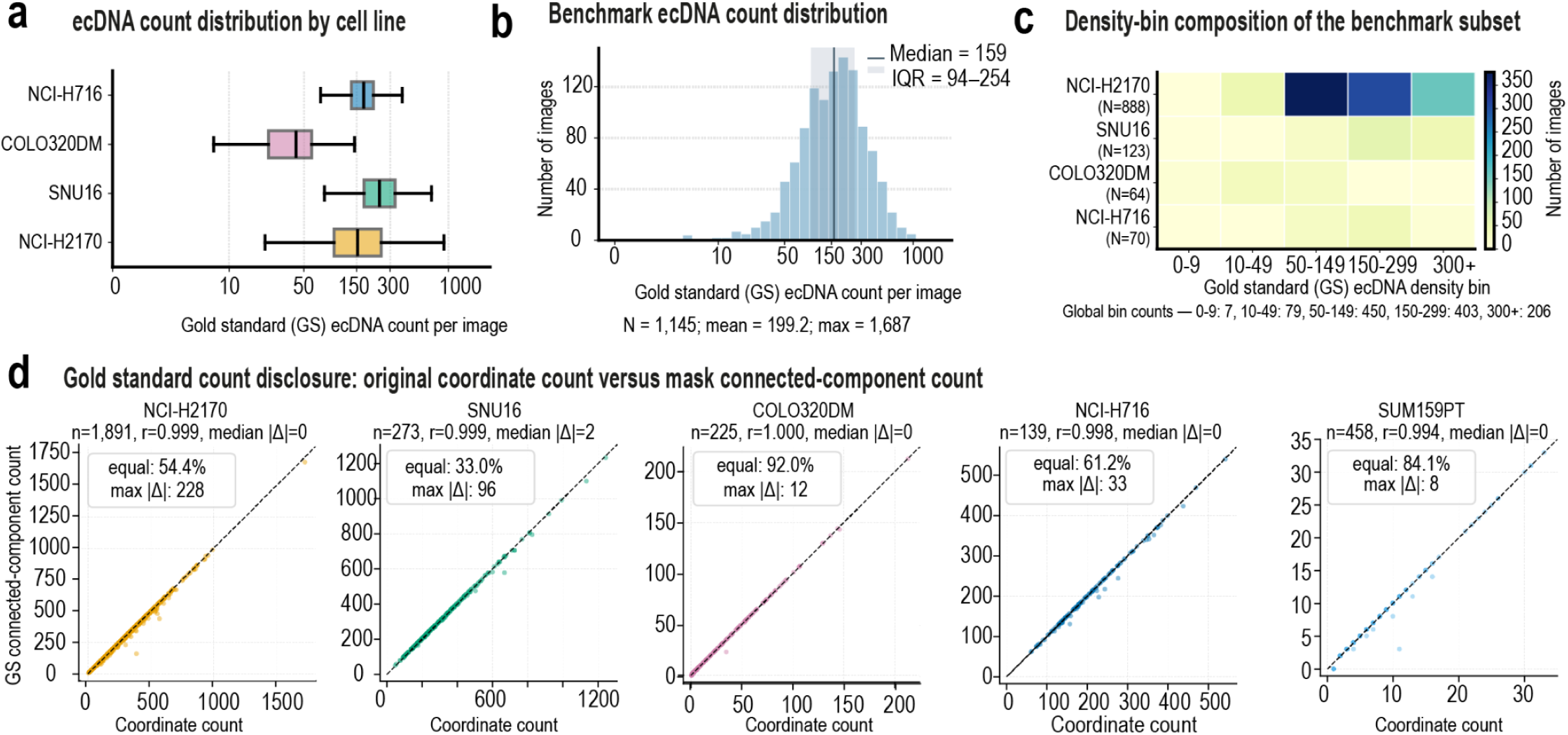
Benchmark composition, ecDNA-density structure and gold-standard count concordance. a, Per-image gold-standard ecDNA abundance across the 1,145-image benchmark, stratified by cell line and shown on a logarithmic scale (NCI-H2170, n = 888; SNU16, n = 123; NCI-H716, n = 70; COLO320DM, n = 64). Boxes indicate the median and interquartile range (IQR); whiskers extend to 1.5 × IQR. **b**, Distribution of gold-standard ecDNA abundance across the benchmark (n = 1,145; median, 159; IQR, 94–254; mean, 199.2; maximum, 1,687), spanning approximately three orders of magnitude in ecDNA burden. **c**, Composition of gold-standard ecDNA-count bins across cell lines. Rows indicate cell lines and columns indicate the five count bins, with each cell reporting the number of images. Global bin counts are 0–9, n = 7; 10–49, n = 79; 50–149, n = 450; 150–299, n = 403; and ≥300, n = 206. NCI-H2170 contributes 888 of the 1,145 benchmark images and predominates in the two central count bins. **d**, Comparison of gold-standard counting representations across the complete 2,986-image resource. For each of five cell lines, raw annotated coordinate counts are compared with 8-connected-component counts derived from rendered gold-standard masks using a minimum component area of 3 pixels. SUM159PT is included as part of the complete imaging resource but is not included in the 1,145-image benchmark. Agreement between representations was high across all cell lines (r ≥ 0.994), with a median absolute difference of 0 for all cell lines except SNU16 (median absolute difference, 2). Differences arise when closely spaced annotation points merge into a single connected component; connected-component counts were used as the canonical benchmark quantity throughout.

**Extended Data Figure 2:**
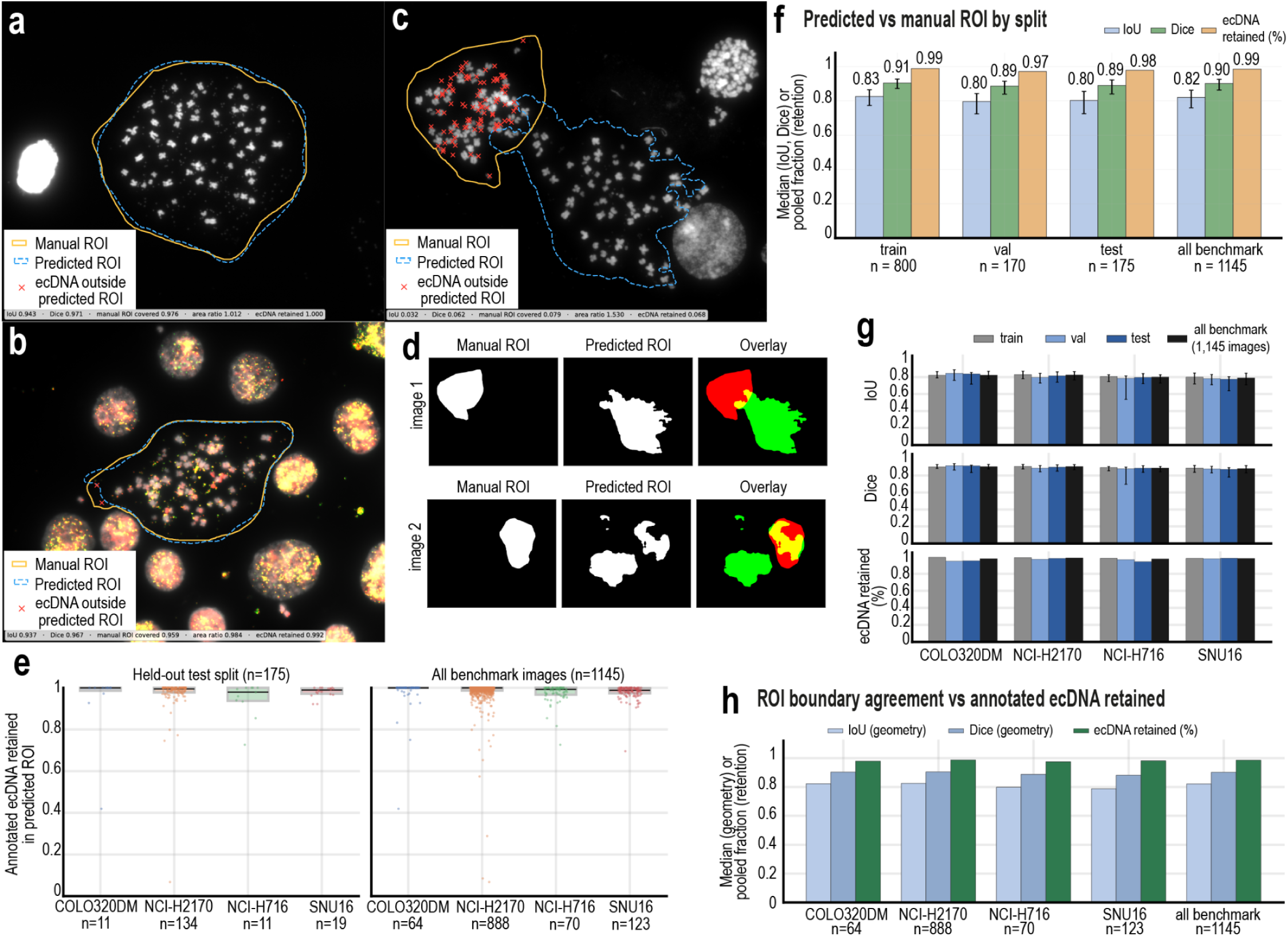
Automated region-of-interest prediction preserves annotated ecDNA despite geometric differences from manual regions. a–c, Representative comparisons of manually defined ROIs (gold) and predicted ROIs (dashed blue), with red crosses indicating annotated ecDNA signals outside the predicted region. High-agreement examples are shown using the grayscale image (a) and RGB composite (b). **c**, Lowest-agreement example in the benchmark, in which the predicted ROI identifies a different metaphase from the manually annotated region. IoU, Dice coefficient and ecDNA retention are shown for each image. **d**, Binary-mask representation of manual and predicted ROIs for two representative images. Green indicates overlap between manual and predicted regions, red indicates manual ROI area excluded by the prediction, and yellow indicates predicted area outside the manual ROI. **e**, Fraction of manually annotated ecDNA signals retained within predicted ROIs, stratified by cell line, for the 175-image held-out test set and complete 1,145-image benchmark. **f**, Median intersection over union (IoU), median Dice coefficient and pooled ecDNA retention across fixed training, validation and held-out test partitions. Comparable performance across partitions indicates that ROI agreement is maintained on images excluded from model development. **g**, IoU, Dice coefficient and ecDNA retention stratified by cell line and dataset partition. **h**, Relationship between geometric ROI agreement and retention of annotated ecDNA across cell lines. Retention of ecDNA exceeds geometric agreement in every cell line because regions of the manual ROI excluded by the prediction contain four- to tenfold fewer annotated ecDNA signals per unit area than regions retained by the predicted ROI, indicating that most boundary disagreement occurs in areas with little annotated ecDNA signal.

**Extended Data Figure 3:**
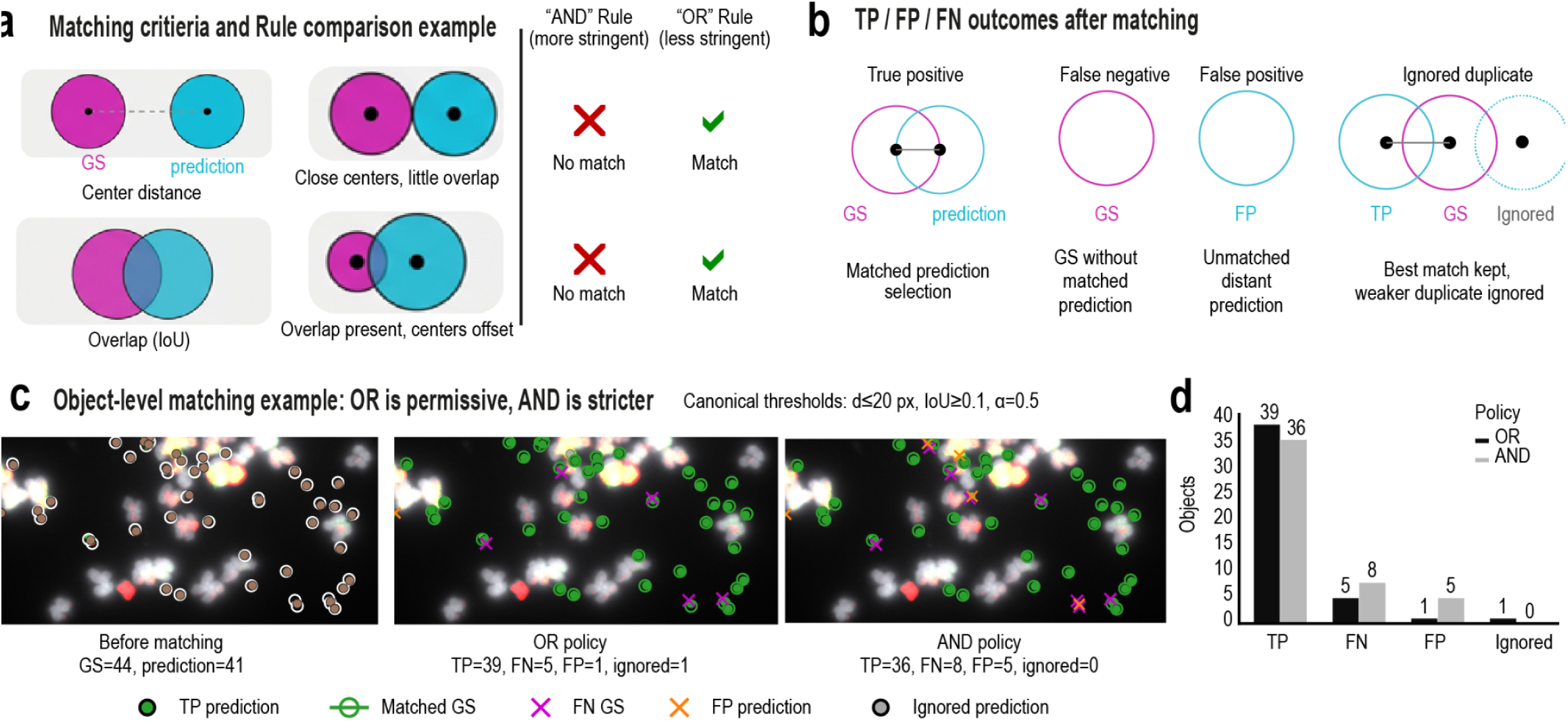
Object-level matching enables tolerance-aware comparison of ecDNA detections. a, Object-level matching criteria based on centroid distance and mask overlap (intersection over union, IoU). Under the AND policy, both criteria must be satisfied for a gold-standard–prediction pair to be eligible for matching; under the OR policy, either criterion is sufficient. The policies therefore differ for detections with close centroids but limited overlap or overlapping masks with offset centroids. **b**, Possible outcomes of one-to-one assignment: true positive, representing a matched gold-standard–prediction pair; false negative, representing a gold-standard object without a matched prediction; false positive, representing a prediction without a valid gold-standard match; and ignored duplicate, representing an additional prediction competing for a gold-standard object assigned to a better candidate. Ignored duplicates are tracked separately and excluded from the false-positive count. **c**, Representative matching outcome for a benchmark image containing 44 gold-standard objects and 41 predictions at the canonical operating point (maximum centroid distance, 20 pixels; minimum IoU, 0.1; α = 0.5). The primary OR policy identified 39 true positives, 5 false negatives, 1 false positive and 1 ignored duplicate. Applying the stricter AND policy to the same image identified 36 true positives, 8 false negatives and 5 false positives, with no ignored duplicates.

**Extended Data Figure 4:**
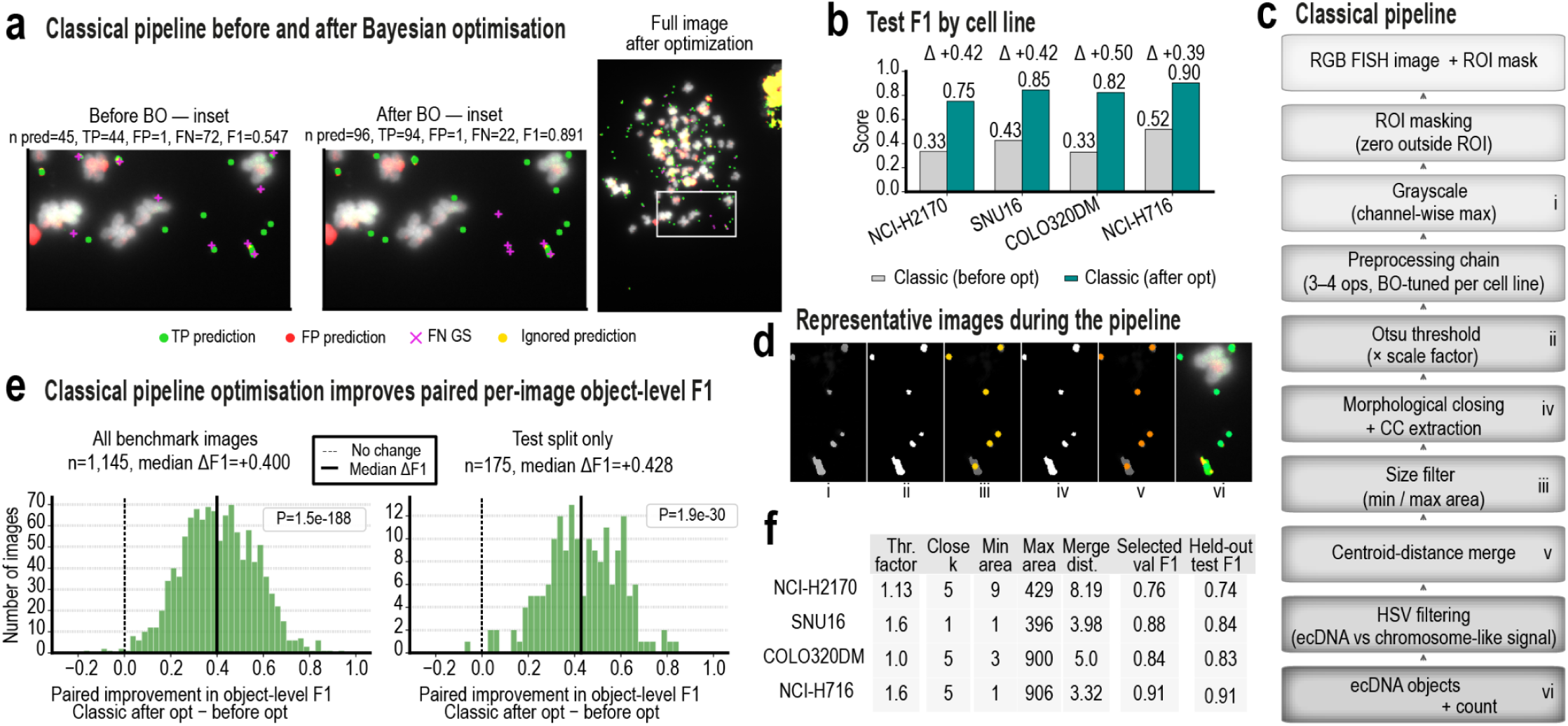
Optimization substantially improves classical ecDNA detection across images and cell lines. a, Representative benchmark metaphase before and after optimization of the classical computer-vision pipeline. Before optimization, the pipeline produced 45 predictions, comprising 44 true positives, 1 false positive and 72 false negatives (F1 = 0.547); after optimization, it produced 96 predictions, comprising 94 true positives, 1 false positive and 22 false negatives (F1 = 0.891). The improvement primarily reflects recovery of previously missed ecDNA objects. **b**, Object-level F1 on the 175-image held-out test set before and after optimization, stratified by cell line and scored with the unified evaluation framework under the canonical OR matching policy. Optimized performance pools to F1 = 0.775 across the held-out test set. **c**, Classical detection workflow comprising ROI masking, channel-wise maximum grayscale conversion, cell-line-specific preprocessing, thresholding at a tuned multiple of the global Otsu threshold, morphological closing, connected-component extraction, area filtering, centroid-distance merging and HSV-based classification of ecDNA-like and chromosome-like signal. Roman numerals correspond to the processing stages shown in d. **d**, Representative image illustrating the sequential processing stages of the classical pipeline. **e**, Paired per-image change in object-level F1 after optimization across the complete 1,145-image benchmark and the 175-image held-out test set. Median ΔF1 was +0.400 across the full benchmark (P = 1.5 × 10⁻¹⁸⁸) and +0.428 on the held-out test set (P = 1.9 × 10⁻³⁰; two-sided Wilcoxon signed-rank test), indicating broad improvement across images rather than gains driven by a limited subset. **f**, Frozen cell-line-specific configurations selected by validation F1, together with the test F1 recorded for each selected configuration at selection time. Selection-time test F1 was computed inside the optimization code and differs slightly from the evaluation-framework values in b, which are used for all comparative analyses.

**Extended Data Figure 5:**
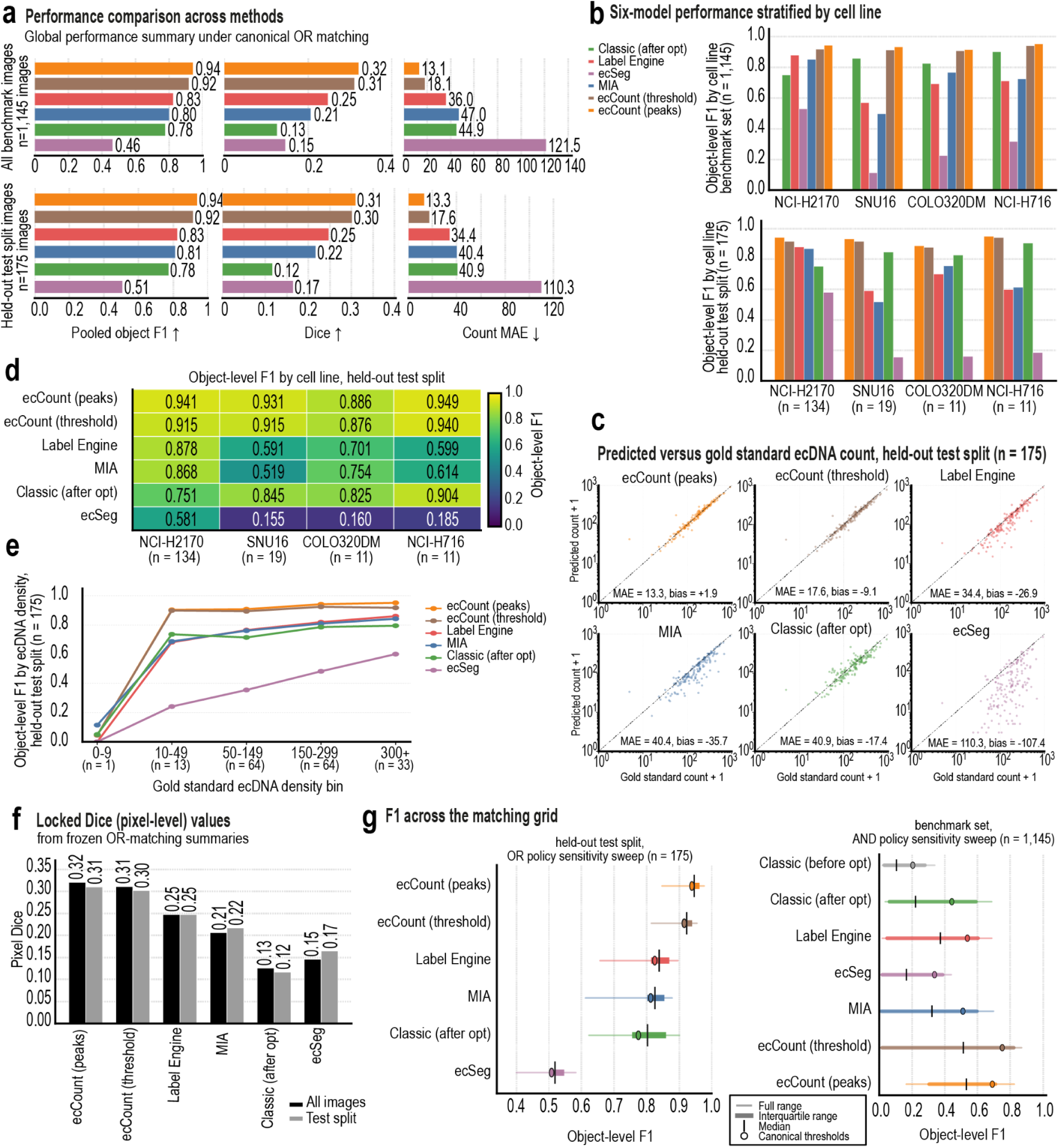
Benchmark performance is preserved on the held-out test set across methods, cell lines and matching criteria. a, Pooled object-level F1, pixel-level Dice and count error for all six evaluated model outputs across the complete 1,145-image benchmark (upper row) and the 175-image held-out test set (lower row). Method ranking is preserved between the full benchmark and held-out test set. **b**, Object-level F1 stratified by cell line for the complete benchmark and held-out test set. **c**, Predicted versus gold-standard ecDNA count on the held-out test set for each method, with count error and mean signed bias shown as insets. ecCount using peak extraction shows substantially lower signed bias relative to its overall count error than the other evaluated approaches. **d**, Object-level F1 by cell line on the held-out test set. **e**, Object-level F1 stratified by gold-standard ecDNA burden on the held-out test set. The 0–9 ecDNA bin contains one image and is shown for completeness. **f**, Pixel-level Dice for all six methods across the complete benchmark and the held-out test set, pooled over all pixels. Values are low for every method because the gold-standard masks are 5 × 5-pixel diamonds placed at annotated points rather than object outlines; object-level metrics are therefore the primary measure. **g**, Object-level F1 over a 10 × 7 grid of matching parameters (maximum centroid distance 5–100 pixels × minimum IoU 0–0.5; 70 combinations). For each method, the thin line spans the range, the bar the interquartile range and the tick the median across the grid; the open circle marks the canonical operating point (20 pixels, 0.1). Left, held-out test set under the OR policy; right, complete benchmark under the AND policy, which also includes the unoptimized classical pipeline. Under OR, the two ecCount outputs rank first and second at every grid point.

**Extended Data Figure 6:**
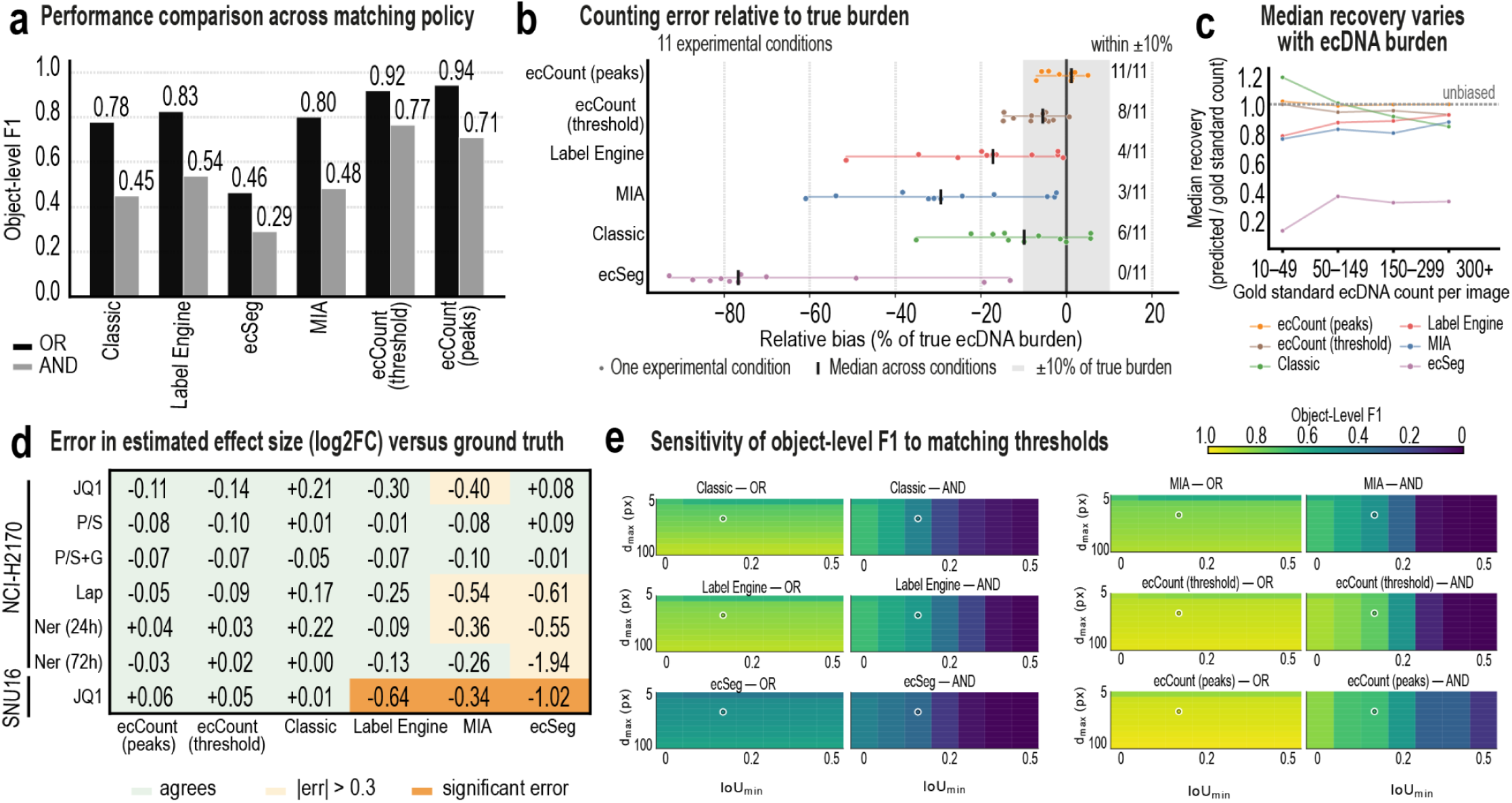
Matching policy and count-dependent error influence quantitative ecDNA conclusions. a, Pooled object-level F1 for all six evaluated model outputs under OR and AND matching policies across the complete 1,145-image benchmark. Under the OR policy, a prediction and gold-standard object are eligible for matching when either the centroid-distance or overlap criterion is satisfied; under the AND policy, both criteria are required. F1 decreases under the stricter AND policy for all methods, while overall method ranking is largely preserved. OR matching is used as the canonical policy throughout the study. **b**, Counting error relative to ecDNA burden across experimental conditions, expressed as a percentage of the corresponding gold-standard count. Control conditions within each cell line are pooled. The shaded region denotes ±10% of gold-standard burden, the vertical marker indicates the median across conditions, and the number of conditions within this range is shown at right. ecCount using peak extraction remains within ±10% for all evaluated conditions. **c**, Median recovery, defined as predicted count divided by gold-standard count, as a function of ecDNA burden. Variation in recovery with burden indicates that counting error changes systematically across the measurement range and can differentially affect experimental comparisons. **d**, Error in estimated log2 fold change relative to gold-standard annotations across seven treated-versus-control comparisons from two cell lines. Each value is the method’s log2 fold change minus the gold-standard log2 fold change. Color indicates agreement; an absolute error above 0.3 with the same significance call; or a different significance call (two-sided Mann–Whitney U, P < 0.05). Both ecCount output modes and the optimized classical pipeline reproduce the gold-standard call in every contrast. SNU16 JQ1 is the only comparison in which significance classification changes for any method and is examined further in main Fig. 6i**. e**, Object-level F1 on the held-out test set over the same 10 × 7 matching grid, one heatmap per method and policy. Minimum IoU is shown on the horizontal axis and maximum centroid distance on the vertical axis; the open circle denotes the canonical operating point. For distance thresholds of 10 pixels or more and minimum IoU of 0.05 or more, F1 depends only on the distance threshold under OR and only on the minimum IoU under AND.

**Extended Data Figure 7:**
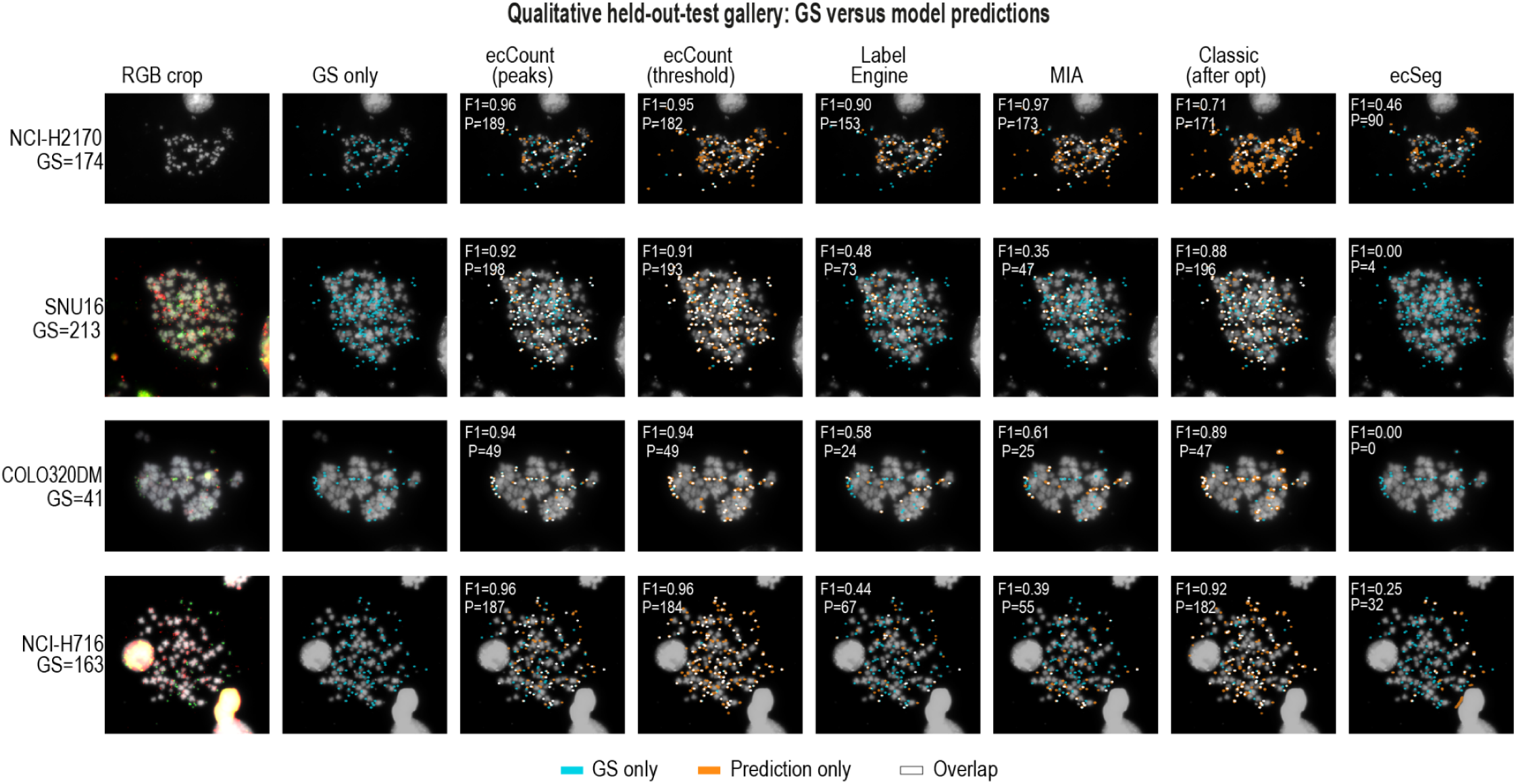
Held-out test images reveal model-specific detection failures across cell lines. Representative held-out metaphase from each benchmarked cell line, selected near that cell line’s median gold-standard ecDNA count: NCI-H2170 (GS = 174), SNU16 (GS = 213), COLO320DM (GS = 41) and NCI-H716 (GS = 163). Columns show the RGB image crop, gold-standard annotation and predictions from each of the six evaluated model outputs. White marks matched detections, cyan marks gold-standard objects without a matched prediction, and orange marks predictions without a matched gold-standard object; per-image object-level F1 and predicted ecDNA count are shown as insets. Matching used the canonical OR policy (maximum centroid distance, 20 pixels; minimum IoU, 0.1; α = 0.5). Label Engine and MIA show substantial under-recovery in SNU16 and NCI-H716, whereas both ecCount output modes retain high object-level agreement. ecSeg produced no detections in the representative SNU16 and COLO320DM images.

**Extended Data Figure 8:**
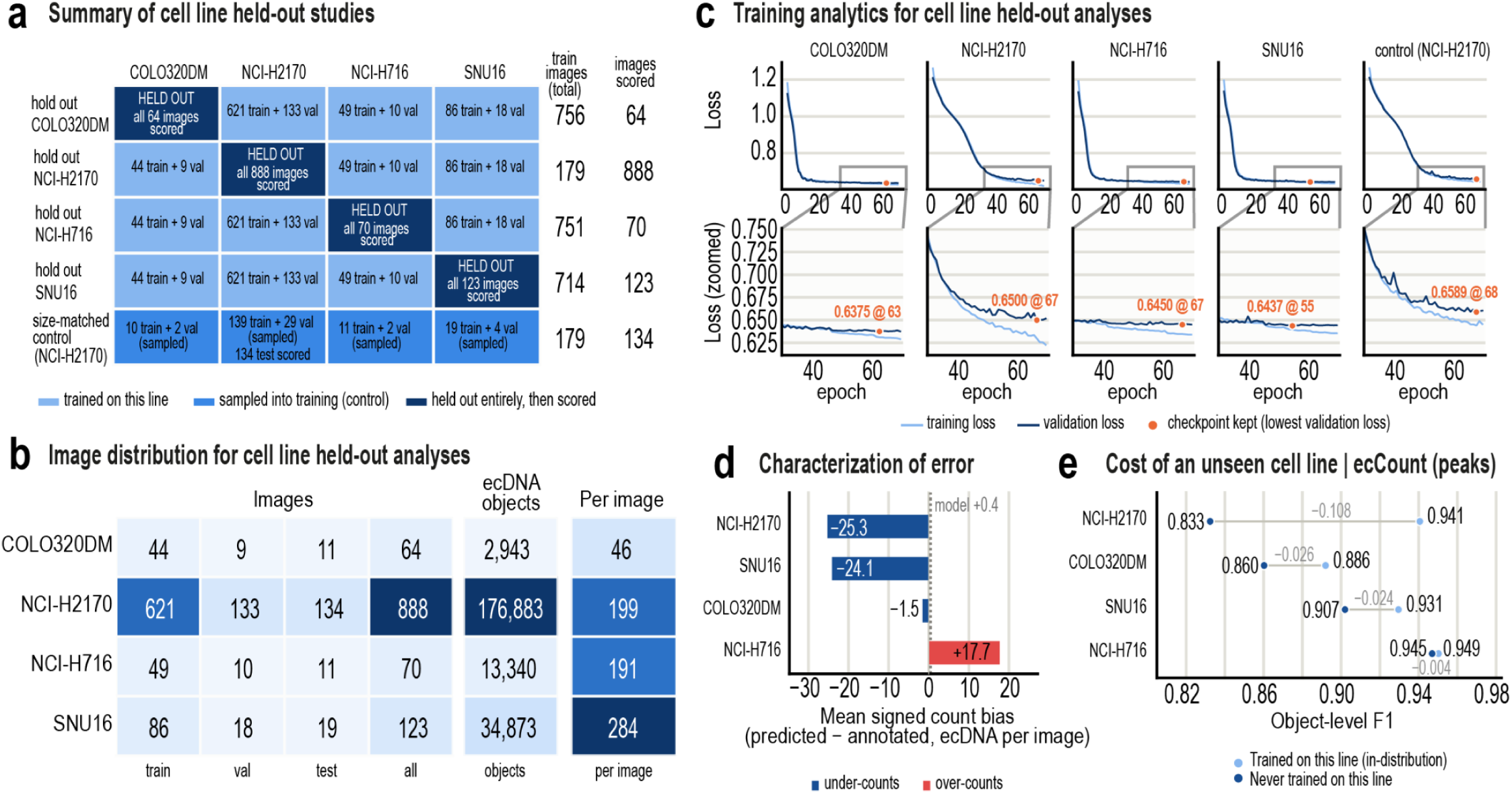
Leave-one-cell-line-out analysis evaluates ecCount transfer to unseen cellular contexts. a, Design of the leave-one-cell-line-out experiments. Rows indicate the cell line excluded from model development and columns indicate the cell lines represented during training or evaluated after training. Color denotes whether a cell line was included in training, sampled as part of the size-matched NCI-H2170 control, or held out entirely and subsequently scored. The total number of training images and images scored is shown for each experiment. Because the size-matched control was exposed to NCI-H2170 during training, it was scored on the NCI-H2170 held-out test images only. **b**, Composition of the 1,145-image benchmark by cell line, showing training, validation, test and total image counts, annotated ecDNA objects and objects per image. NCI-H2170 contributes 78% of the benchmark images and 78% of its annotated ecDNA objects. **c**, Training and validation loss trajectories for each leave-one-cell-line-out model and the size-matched NCI-H2170 control. Enlarged views show the region surrounding the retained checkpoint, selected by minimum validation loss. Validation sets differ between runs, so these losses index checkpoint selection within a run and are not comparable across panels. **d**, Mean signed count bias for ecCount peak detection on each unseen cell line, calculated as predicted minus annotated ecDNA count per image. Negative values indicate systematic undercounting and positive values indicate overcounting; the dashed line marks the released model’s bias of +0.4 ecDNA per image across the complete benchmark. **e**, Cost of cellular-context novelty for ecCount peak detection. Object-level F1 for the model trained without each cell line is compared with the released model, both scored on the held-out test images of that cell line; connecting lines show the difference, computed at full precision before rounding. The same four values are shown as bars in Fig. 6f. Three of the four comparisons rest on fewer than 20 images. OR matching (maximum centroid distance, 20 pixels; minimum IoU, 0.1; α = 0.5).

**Extended Data Figure 9:**
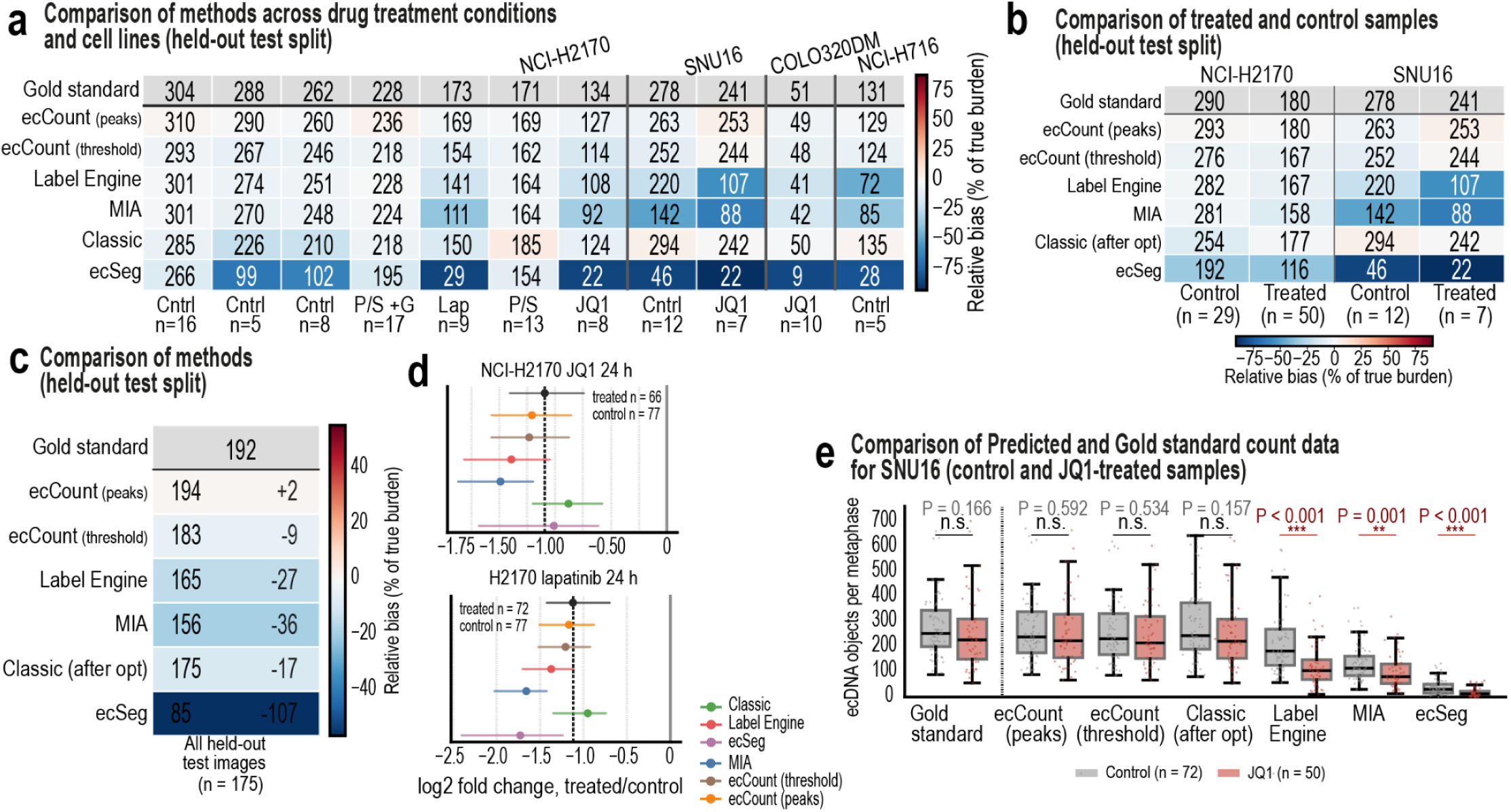
Treatment analyses confirm that counting bias can alter inference of ecDNA dynamics. a, Comparison of computational methods across drug-treatment conditions and cell lines using only images from the 175-image held-out test split. Rows indicate methods and columns indicate experimental conditions. Values show predicted ecDNA count per image, and cell color denotes relative bias from the corresponding gold-standard burden (blue, undercounting; red, overcounting; white, minimal bias). Gold-standard mean counts are shown for each condition. **b**, Comparison of treated and control populations for held-out NCI-H2170 and SNU16 images. Treatment conditions were pooled within each cell line to increase the number of held-out images available for comparison. Values indicate mean ecDNA counts and cell color denotes relative bias from gold standard as in ***a***. **c**, Aggregate performance across all 175 held-out test images, combining treatment conditions and cell lines. Mean predicted ecDNA counts are shown for each method alongside the corresponding gold-standard count, with relative bias indicated by cell color. **d**, Estimated treatment effects for two NCI-H2170 perturbation comparisons across the complete benchmark. Points indicate log2 fold change in ecDNA burden for treated versus control cells, with gold-standard effect sizes indicated by dashed vertical lines. Differences between predicted and gold-standard effect sizes illustrate how count-dependent bias can distort the magnitude, and in some cases the interpretation, of treatment-associated changes; ecCount peak detection remains closest to the gold-standard effect across comparisons. **e**, Distribution of predicted ecDNA counts for control and JQ1-treated SNU16 cells at 24 hours across the complete benchmark, shown per method as the distribution underlying the effect sizes in Fig. 6i. Manual annotations show no significant difference between conditions, a conclusion reproduced by ecCount peak detection, ecCount threshold-mask extraction and the optimized classical pipeline. Label Engine, MIA and ecSeg instead predict significant differences between control and treated populations, illustrating how systematic counting bias can alter the inferred biological response.

